# The histone chaperone FACT induces Cas9 multi-turnover behavior and modifies genome manipulation in human cells

**DOI:** 10.1101/705657

**Authors:** Alan S. Wang, Leo Chen, R. Alex Wu, Christopher D. Richardson, Benjamin G. Gowen, Katelynn R. Kazane, Jonathan T. Vu, Stacia K. Wyman, Jiyung Shin, Johannes C. Walter, Jacob E. Corn

## Abstract

Cas9 is a prokaryotic RNA-guided DNA endonuclease that binds substrates tightly *in vitro* but turns over rapidly when used to manipulate genomes in eukaryotic cells. Little is known about the factors responsible for dislodging Cas9 or how they influence genome engineering. Using a proximity labeling system for unbiased detection of transient protein interactions in cell-free *Xenopus laevis* egg extract, we identified the dimeric histone chaperone FACT as an interactor of substrate-bound Cas9. Immunodepletion of FACT subunits from extract potently inhibits Cas9 unloading and converts Cas9’s activity from multi-turnover to single-turnover. In human cells, depletion of FACT delays genome editing and alters the balance between indel formation and homology directed repair. Depletion of FACT also increases epigenetic marking by dCas9-based transcriptional effectors with concomitant enhancement of transcriptional modulation. FACT thus shapes the intrinsic cellular response to Cas9-based genome manipulation most likely by determining Cas9 residence times.

## Introduction

Cas9 is a Clustered Regularly Interspaced Short Palindromic Repeats (CRISPR)-associated RNA-guided DNA endonuclease that is directed to a target DNA molecule by forming a ribonucleoprotein (RNP) complex with a guide RNA (gRNA) (Doudna and Charpentier, 2014; Hsu et al., 2014; Jinek et al., 2012; Knott and Doudna, 2018). After unwinding its duplex substrate, Cas9 uses two nuclease domains to generate a double-stranded break (DSB) (Doudna and Charpentier, 2014; Hsu et al., 2014; Jinek et al., 2012; Knott and Doudna, 2018; Sternberg et al., 2015). Subsequent break repair relies on a cell’s endogenous machinery to either incorporate sequences using a DNA template through homology-directed repair (HDR) or introduce insertions or deletions (indels) during non-homologous end joining (NHEJ) (Maggio and Goncalves, 2015). The ease of programming Cas9 to generate targeted DSBs and initiate break repair has enabled its widespread use as a genome editing agent (Cong et al., 2013; Jinek et al., 2012; 2013; Mali et al., 2013).

Mutations that inactivate Cas9’s nuclease activity preserve its capacity to bind a gRNA and target DNA (Qi et al., 2013). Catalytically inactive Cas9 (dCas9) has dramatically expanded the CRISPR toolbox. Fusing various effector proteins to dCas9 has enabled CRISPR-based methods to activate or repress gene expression, manipulate the three-dimensional architecture of nuclei, image genomic loci, track RNA molecules, and identify proteins at specific loci (Chen et al., 2013; Gao et al., 2018; Gilbert et al., 2013; Hilton et al., 2015; Konermann et al., 2015; Liu et al., 2017; Myers et al., 2018; Nelles et al., 2016; Qi et al., 2013; Schmidtmann et al., 2016; Wang et al., 2018).

The relationship between the efficacy of the CRISPR-Cas9 toolbox and Cas9’s lifetime on a genomic target site is unclear. For example, CRISPR transcriptional effectors localize histone acetyltransferases or methyltransferases around a transcription start site (TSS) to manipulate expression of endogenous genes (Gilbert et al., 2013; Hilton et al., 2015). Such transcriptional engineering presumably relies on providing these histone modifiers sufficient time at the TSS to deposit the appropriate epigenetic marks. Conversely, Cas9’s utility as a targeted nuclease may be predicated on its removal from the genome because Cas9 itself masks the DSB from cellular repair enzymes (Clarke et al., 2018; Richardson et al., 2016b). Cas9 residence times and unloading might thus play crucial roles in Cas9-based interventions.

While some Cas9 molecules behave as multi-turnover enzymes (Yourik et al., 2019), the widely used *Streptococcus pyogenes* Cas9 and dCas9 exhibit extremely stable protein-DNA interactions and possess residence times of over five hours *in vitro* (Raper et al., 2018; Richardson et al., 2016b; Sternberg et al., 2014). Estimates of residence times in live cells vary (Ma et al., 2016; Shao et al., 2016), but some experiments indicate that *S. pyogenes* Cas9 stays bound to its target in mammalian cells for as little as five minutes (Knight et al., 2015) and imply that cellular factors promote turnover. The ability to detect resolved genomic edits only a few hours after electroporation of Cas9 RNPs (Kim et al., 2014) further suggests that cells actively remove Cas9 from the genome either purposefully or as a byproduct of normal genome metabolism.

Prior work has suggested that RNA polymerases can dislodge Cas9 from DNA *in vitro* when the gRNA anneals to the non-coding DNA strand but not to the coding strand (Clarke et al., 2018). Targeting the non-coding strand with a gRNA roughly correlated with increased editing rates in human cells and an increased ability of a bacterially-encoded CRISPR system to fight phage infection. However, the ability to edit non-transcribed regions of the human genome implies that RNA polymerases are not solely responsible for Cas9 eviction. Moreover, the ability to edit post-mitotic cells such as neurons suggests that replicative DNA polymerases are also not solely responsible for unloading Cas9 (Nishiyama et al., 2017; Suzuki et al., 2016).

Here, we find that metazoan cellular extracts contain a factor responsible for rapid multi-turnover activity of Cas9 on DNA substrates. An unbiased proteomics approach to mark proteins transiently associating with substrate-bound Cas9 and dCas9 identified both components of the heterodimeric Facilitates Chromatin Transcription (FACT) histone chaperone complex, SPT16 and SSRP1. Immunodepletion of FACT subunits in extract was sufficient to prevent dCas9 removal and converted Cas9’s activity from multi-turnover to single-turnover. In living human cells, FACT modified Cas9 editing outcomes and played a strand-independent role in determining the extent of epigenetic marking and transcriptional regulation from dCas9-based effectors. These results reveal an unanticipated functional interaction between Cas9 and the eukaryotic machinery responsible for regulating nucleosome assembly. Manipulating FACT provides a means to modify dCas9 residence times and thereby improve the efficacy of the CRISPR-Cas9 toolbox.

## Results

### Cell-Free *X. laevis* Egg Extract Promotes Rapid Turnover of Cas9 from DNA Substrates

*Xenopus* egg extracts have a long track record of dissecting nuclear dynamics and interrogating processes such as DNA replication, chromosome segregation, and DNA repair (Heald et al., 1996; Hoogenboom et al., 2017; Kalab et al., 2006; Knipscheer et al., 2009; Lebofsky et al., 2009). We therefore used high-speed supernatant (HSS) of total *Xenopus laevis* egg lysate (Lebofsky et al., 2009) to look for cellular factors that promote the dissociation of a pre-formed Cas9 RNP-DNA complex.

We first tested the ability of HSS to promote the dissociation of *S. pyogenes* Cas9 RNPs from linear and plasmid DNA substrates harboring a single on-target site. We incubated RNPs with twice as many moles of DNA for 45 minutes before adding buffer or HSS. With excess substrate, complete cleavage would occur only if Cas9 possesses multi-turnover activity. Consistent with prior *in vitro* data (Richardson et al., 2016b; Sternberg et al., 2014), a Cas9 RNP targeting linear double-stranded DNA in buffer failed to cleave the excess substrate and thus behaved as a single-turnover enzyme even under these multi-turnover conditions (Figure 1A). However, incubation of RNP and target DNA in HSS and ATP yielded a steady conversion of substrate to product that was consistent with multi-turnover behavior (Figure 1A). Pre-depleting HSS ATP levels using calf intestinal phosphatase (CIP) and adding an excess of the non-hydrolysable analogue ATPγS reverted Cas9 to single-turnover behavior (Figure 1A). We observed similar results when targeting Cas9 to a circular plasmid under multi-turnover conditions (Figure 1B).

**Figure 1.**
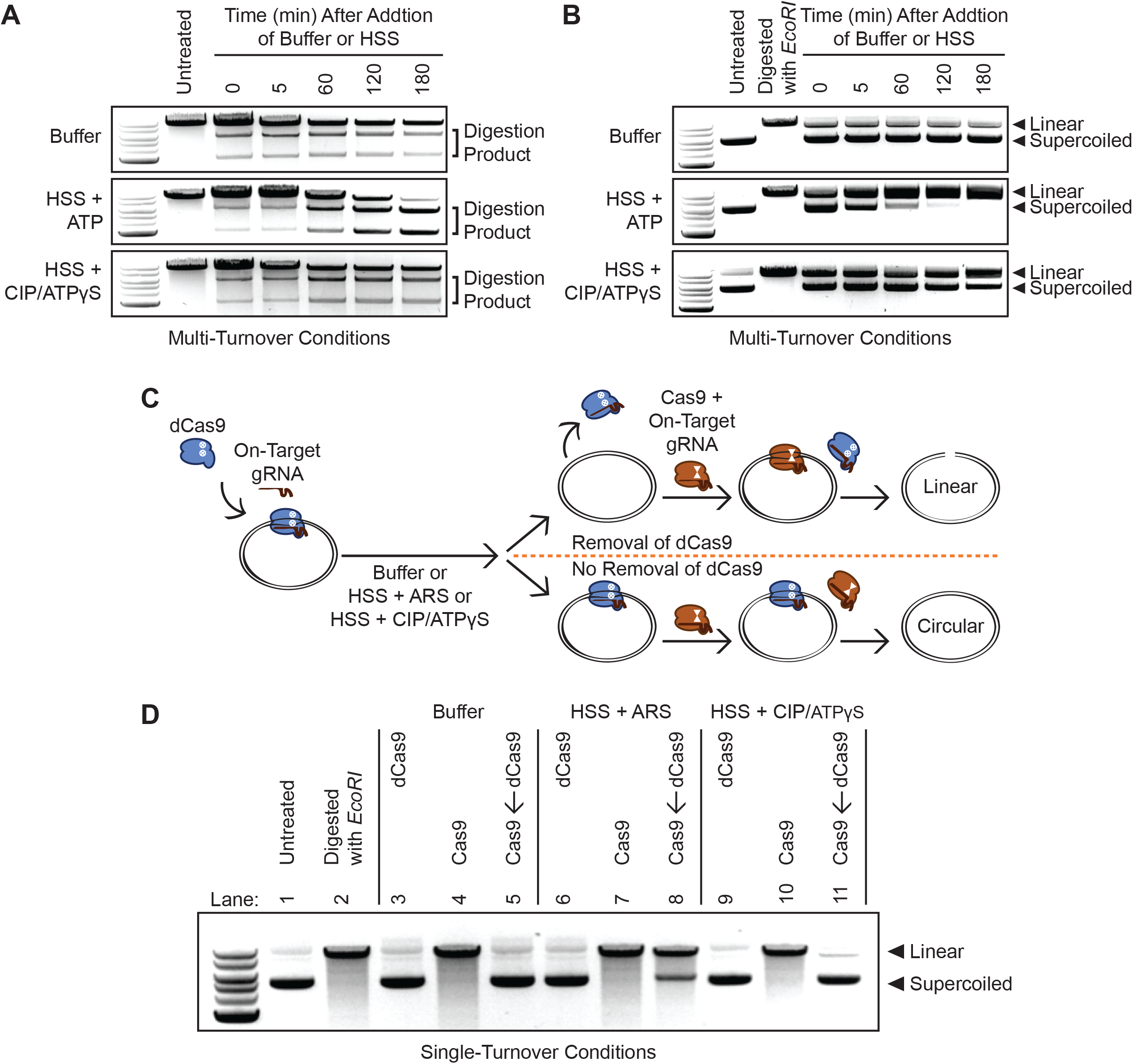
Energy-Dependent Release of Cas9 and dCas9 from DNA in HSS. (A) Time course of Cas9 RNPs programmed against a linear DNA substrate in a 1:2 molar ratio in buffer, HSS with ATP, or HSS with CIP and ATPγS. (B) Time course of Cas9 RNPs programmed against a plasmid substrate in a 1:2 molar ratio in buffer, HSS with ATP, or HSS with CIP and ATPγS. (C) Schematic of the single-turnover competition assay. (D) ATP-dependent unloading of dCas9 off a plasmid substrate in *X. laevis* HSS. Presence of linearized DNA after addition of a 10-fold molar excess of Cas9 indicates removal of dCas9 while persistence of circular DNA indicates stable binding of dCas9.

We next developed a competition assay to determine whether factors in HSS induce dCas9 removal from a circular DNA substrate under single turnover conditions (Figure 1C). We allowed a dCas9 RNP to equilibrate in buffer with a plasmid containing a single on-target site and then incubated this RNP-plasmid complex with buffer alone, HSS containing an ATP-regenerating system (ARS, see Methods), or HSS pre-incubated with CIP and excess ATPγS. Finally, we added a 10-fold excess of catalytically active Cas9 programmed with the identical on-target gRNA. Persistent binding of dCas9 should prevent binding of Cas9 and preclude cleavage while dCas9 unloading would grant Cas9 access to the target site. The presence of linearized DNA thus provides a readout of dCas9 unloading. Consistent with our data using multi-turnover conditions, buffer alone did not promote dCas9 dissociation from the plasmid (Figure 1D Lanes 3-5). By contrast, HSS containing the ARS rapidly removed dCas9 to allow the catalytically active Cas9 to cleave the plasmid within 45 minutes (Figure 1D Lanes 6-8). Conversely, ATP-depleted HSS supplemented with ATPγS did not evict the majority of dCas9 molecules, largely preventing Cas9 from cleaving the plasmid (Figure 1D Lanes 9-11). Overall, our data with linear and circular DNA substrates indicate that HSS contains at least one factor capable of dislodging Cas9 from its target, enabling re-binding to uncleaved molecules and multi-turnover behavior.

### Unbiased Cas9 Interaction Marking Identifies the FACT Histone Chaperone as Required for Cas9 Removal and Multi-Turnover Activity in HSS

To identify factors that remove Cas9 from DNA in HSS, we fused Cas9 and dCas9 to the promiscuous biotin ligase BirA* (Arg118Gly). BirA* covalently labels nearby proteins, which can then be isolated with Streptavidin-coupled beads and identified through mass spectrometry proteomics (Gao et al., 2018; Roux et al., 2012; Schmidtmann et al., 2016). BirA*’s utility for our purposes is derived from our ability to extend the labeling time beyond a few minutes, thereby allowing the system to accumulate biotinylated versions of transient yet repeated interactors (Roux et al., 2013). BirA*-dCas9 fusions expressed in living cells have helped identify DNA interactors at repetitive genomic regions (Schmidtmann et al., 2016), but excess BirA*-dCas9 unbound to the genomic target has complicated its use at non-repetitive loci. We reasoned that the ability to form a defined Cas9-BirA* RNP-DNA or dCas9-BirA* RNP-DNA species in HSS could enable identification of Cas9 and dCas9 removal factors.

We expressed and purified recombinant Cas9-BirA* and dCas9-BirA* in and from bacterial cells (Data S1 and Supplemental Figure 1A). The fused biotin ligase neither compromised Cas9’s nuclease activity nor hindered rapid dislodging of dCas9 in HSS (Supplemental Figures 1B and 1C). We programmed Cas9-BirA* with an on-target gRNA, dCas9-BirA* with the same on-target gRNA, or dCas9-BirA* with a non-targeting (NT) gRNA. We added a ten-fold molar excess of plasmid substrate relative to each RNP in buffer and then added this mixture to HSS containing ARS and biotin (Figure 2A). Streptavidin pulldown and label-free proteomic mass spectrometry (Wuhr et al., 2014) identified biotinylated *X. laevis* proteins that were specifically enriched by gRNA-mediated binding of Cas9 or dCas9 to the plasmid relative to non-specific biotinylation when dCas9 was complexed with the NT gRNA. Cas9-BirA* and dCas9-BirA* programmed with the on-target gRNA had nearly identical interactors (Supplemental Figure 2A), consistent with prior *in vitro* data indicating that Cas9 obscures a DSB so that repair factors are not preferentially enriched around Cas9 (Clarke et al., 2018; Richardson et al., 2016b).

**Figure 2.**
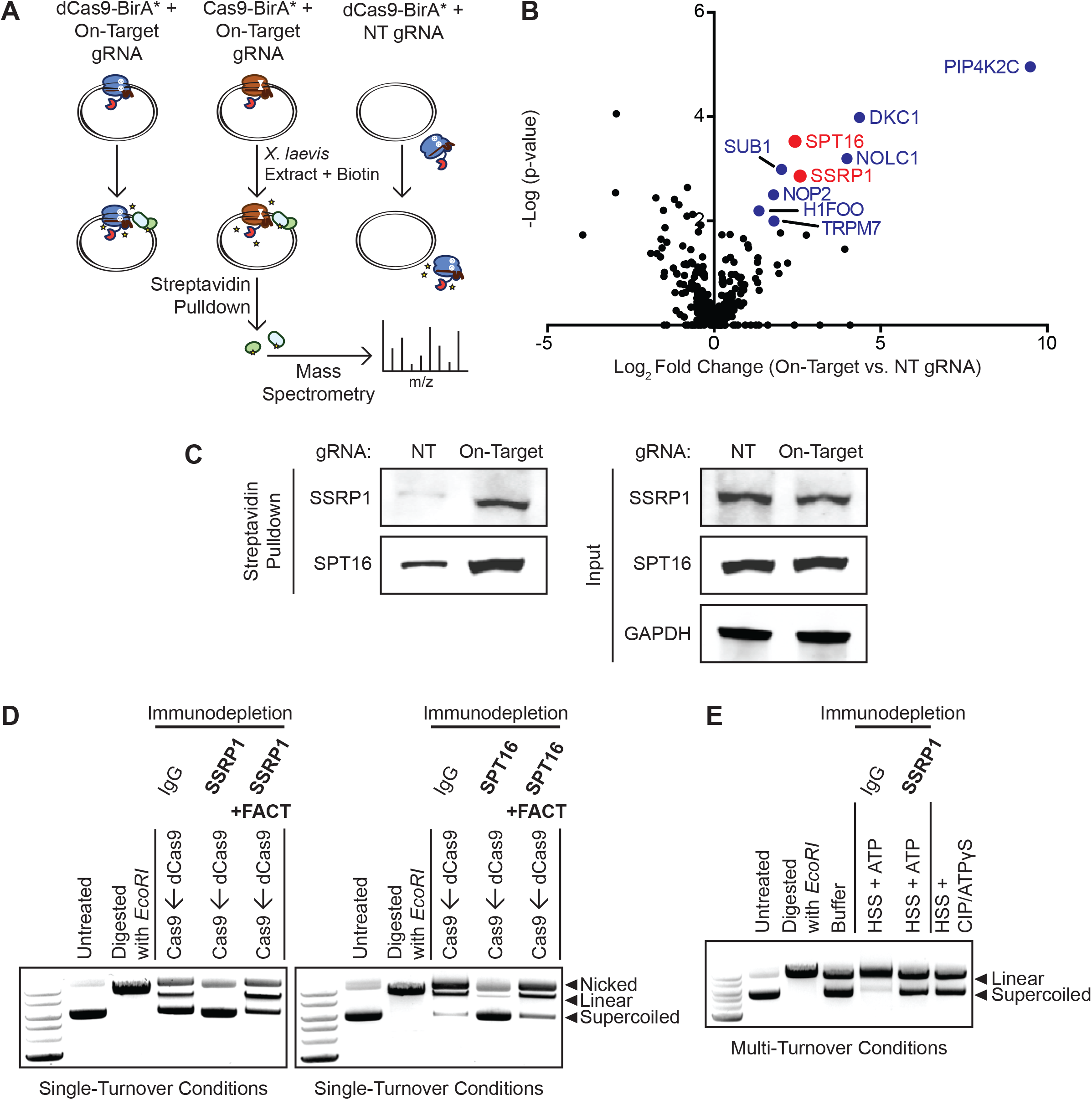
FACT Complex Interacts with DNA-Bound Cas9 and dCas9 to Promote Eviction and Multi-Turnover Behavior. (A) Schematic of samples prepared for mass spectrometry. dCas9-BirA* programmed with the on-target gRNA, Cas9-BirA* programmed with the on-target guide, and dCas9-BirA* programmed with a NT sgRNA were incubated with a 10-fold molar excess of plasmid and then added to HSS containing the ARS and biotin. Biotinylated proteins were isolated with Streptavidin-coupled beads and identified with mass spectrometry. (B) Volcano plot of biotinylated proteins in Cas9-BirA*–on-target gRNA samples (N = 3 biological replicates) versus dCas9-BirA*–NT gRNA samples (N = 3 biological replicates). Colored circles correspond to factors that were significantly enriched (p < 0.05) according to a Limma analysis. Red circles correspond to the two components of the FACT complex. (C) Enrichment of biotinylated SSRP1 and SPT16 in HSS containing dCas9-BirA*–on-target gRNA versus dCas9-BirA*–NT gRNA. (D) FACT immunodepletion inhibits dCas9 eviction in HSS. Addition of a 10-fold molar excess of Cas9 did not generate linearized DNA when dCas9 RNP-plasmid complexes were incubated in SSRP1 or SPT16-immunodepleted extract. Addition of recombinant FACT to SSRP1 or SPT16-immunodepleted extract rescued unloading of dCas9. (E) SSRP1 promotes Cas9’s multi-turnover activity. Cas9 RNPs were incubated with plasmid substrate in a 1:2 molar ratio with either buffer, mock-immunodepleted HSS with ATP, SSRP1-immunodepleted HSS with ATP, or HSS with CIP and ATPγS for 180 min.

Three major sets of DNA-bound Cas9 interactors were apparent by comparing the on-target samples to the NT background control: PIP4K2C; H/ACA-associated proteins DKC1, NHP2, NOP10, and GAR1; and both components of the FACT heterodimer, SPT16 and SSRP1 (Figure 2B and Supplemental Table 1). PIP4K2C is a lipid kinase that converts phosphatidylinositol-4-phosphate to phosphatidylinositol-4,5-bisphosphate. PIP4K2C is not explicitly linked to DNA metabolism, but it has recently been found that phosphoinositides accumulate at sites of double-stranded DNA damage (Wang et al., 2017). H/ACA RNPs are involved in pseudouridylation of RNA, maintenance of telomere integrity, and ribosome biogenesis (Kiss et al., 2010). The ability of H/ACA-associated proteins to interact with unique RNA secondary structures and preserve genomic integrity could imply roles in the cellular response to Cas9 binding. However, PIP4K2C and H/ACA proteins are not known to destabilize protein-DNA interactions and thus were not top candidates for being the Cas9 release factors in HSS.

The FACT complex is a histone chaperone with established roles in nucleosome assembly and remodeling and thus represented an attractive candidate to mediate Cas9 removal in HSS. FACT is a heterodimer consisting of SPT16 and SSRP1, both of which were strongly enriched by proximity biotinylation and unbiased proteomics. FACT promotes chromosomal transactions by removing the H2A-H2B dimer specifically or generally weakening histone contacts within chromatin (Okuhara et al., 1999; Orphanides et al., 1998; Winkler and Luger, 2011). Individual testing by Streptavidin pulldown and Western blotting confirmed that binding of dCas9-BirA* to a plasmid leads to increased biotinylation of SSRP1 and SPT16 (Figure 2C).

To determine whether FACT is responsible for Cas9 removal, we immunodepleted either SSRP1 or SPT16 from HSS (Supplemental Figure 2B). In the single-turnover competition assay, immunodepletion of either FACT component was sufficient to prevent even a 10-fold excess of Cas9 from accessing a DNA target that was pre-bound by dCas9 (Figure 2D). Add-back of recombinant human FACT to SSRP1 or SPT16-immunodepleted extracts rescued the ability of HSS to dislodge dCas9 (Figure 2D). Notably, under multi-turnover conditions, we found that immunodepletion of SSRP1 was sufficient to convert Cas9’s activity from multi-turnover to apparently single-turnover (Figure 2E).

### Knockdown of FACT Alters Cas9 Genome Editing Outcomes in Human Cells

We next asked whether FACT influences Cas9-based interventions in intact human cells. We first examined FACT’s role in shaping genome editing outcomes by measuring editing rates and outcomes via amplicon next generation sequencing (amplicon-NGS) after FACT knockdown (Supplemental Tables 2-4). Transfection of siRNAs to deplete SPT16 led to a concomitant reduction in SSRP1 levels (Supplemental Figure 3A) consistent with prior reports that levels of the two FACT subunits are interdependent (Safina et al., 2013). Sixty hours after transfecting K562 cells with either NT or SPT16 siRNAs, we electroporated separate cultures of cells with Cas9 RNPs targeted to eight different loci, including a non-transcribed gene desert. For each locus, we performed editing reactions with and without a matched single-stranded oligodeoxynucleotide (ssODN) HDR donor that programs a PAM mutation at the appropriate locus.

Knockdown of FACT did not significantly alter indel frequencies measured after 48 hours in the absence of an HDR donor (Supplemental Figure 3B). Consistent with previous reports (Richardson et al., 2016a), inclusion of an ssODN donor increased total editing (indels plus HDR) (Supplemental Figure 3C). This increase in editing was consistent across all eight gRNAs tested and rescued otherwise relatively ineffective gRNAs.

In the presence an ssODN, siRNA-mediated depletion of SPT16 did not affect total editing frequencies relative to the NT siRNA control 48 hours after Cas9 RNP electroporation (Supplemental Figure 3D). However, a time course of editing rates at one site (*VEGFA*) (Supplemental Figure 3E) revealed that SPT16 knockdown significantly impeded the rate of HDR (Figure 3A). HDR levels for SPT16-depleted cells was approximately 50% of that in NT siRNA-treated cells 12 hours after electroporation of Cas9 RNPs (Figure 3A). Indels in SPT16-depleted cells were somewhat reduced at early timepoints but caught up to that of NT siRNA-treated cells within 24 hours (Figure 3B). Measurements taken 48 hours after electroporation indicate that SPT16 knockdown ultimately increased indel frequencies and concomitantly reduced HDR frequencies by up to 50% at multiple loci (Figures 3C and 3D). At the eight loci tested, we observed no marked difference in the effect of SPT16 knockdown on gRNAs that targeted the coding or non-coding strand (Figure 3C and 3D). While knockdown of SPT16 did not alter the relative abundance of the six most common edited alleles for a *CD59* locus, depletion of FACT altered their absolute frequencies and the relative abundance of several alleles each comprising less than 1% of all editing outcomes (Figure 3E). We found similar conservation of the most frequent allele but re-ordering of minor alleles after editing the *VEGFA* locus (Supplemental Figure 3F).

**Figure 3.**
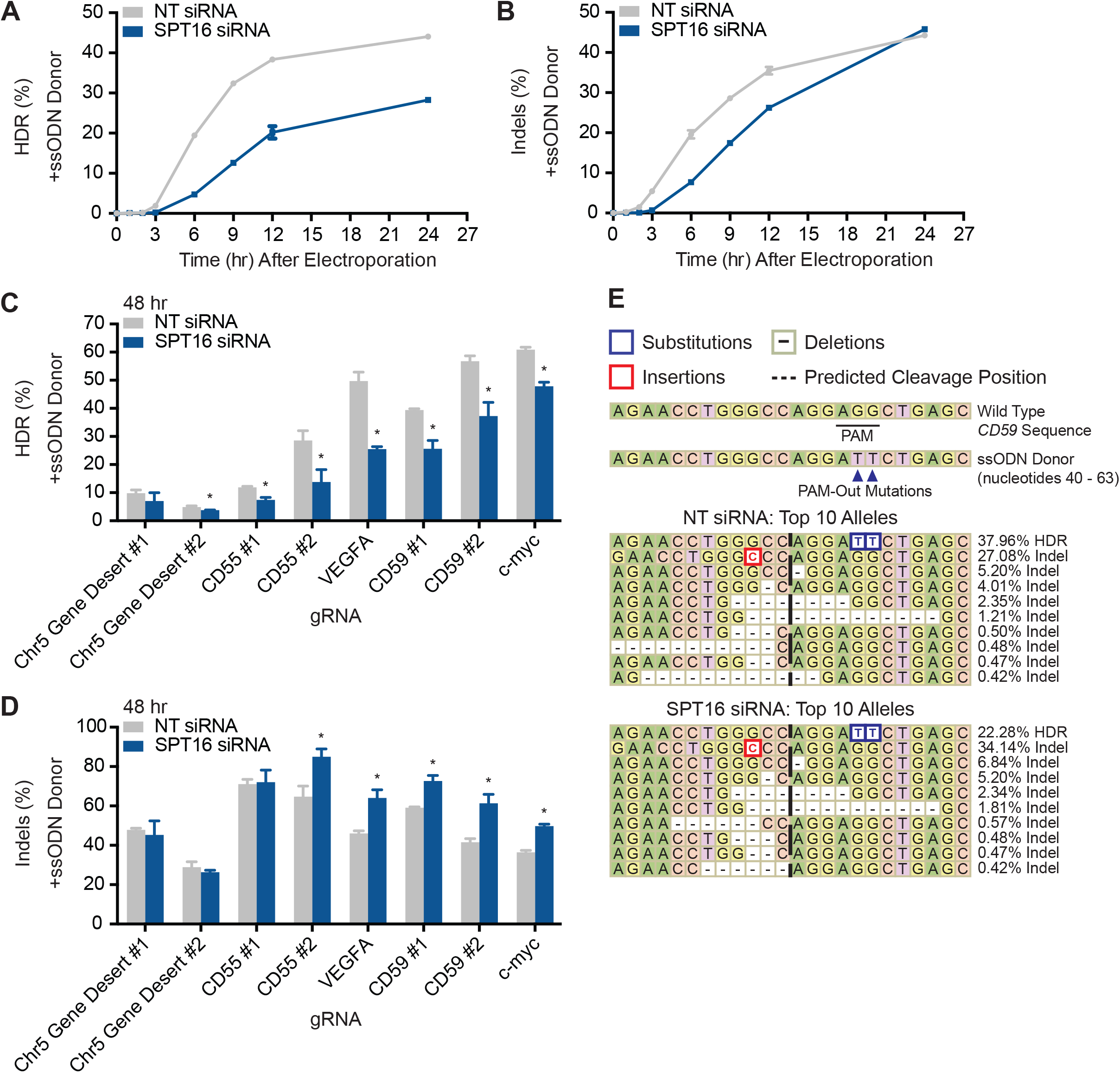
FACT Alters Cas9 Genome Editing Outcomes in Human Cells. (A) HDR rates from amplicon-NGS sequencing of *VEGFA* 0, 1, 2, 3, 6, 9, 12, and 24 hours after electroporation of Cas9 RNPs in the presence of an HDR donor (N = 3 biological replicates). (B) Indel rates from amplicon-NGS sequencing of *VEGFA* 0, 1, 2, 3, 6, 9, 12, and 24 hours after electroporation of Cas9 RNPs in the presence of an HDR donor (N = 3 biological replicates). (C) HDR rates from amplicon-NGS sequencing of eight different loci 48 hours after electroporation of Cas9 RNPs in the presence of an HDR donor (N = 3 biological replicates). (D) Indel rates from amplicon-NGS sequencing of eight different loci 48 hours after electroporation of Cas9 RNPs in the presence of an HDR donor (N = 3 biological replicates). (E) Representative alleles from cells edited with Cas9 programmed with a *CD59* gRNA and a PAM-out ssODN HDR donor.

### Knockdown of FACT Increases Epigenetic Marking and Transcriptional Phenotypes from dCas9-Based Effectors in Human Cells

dCas9-based transcriptional effectors rely on the recruitment of epigenetic modifying enzymes to a target site. We asked whether FACT influences the deposition of chromatin marks by these tools. We examined two different dCas9-based modifiers. The histone acetyltransferase p300 directly deposits H3K27ac marks on chromatin to upregulate transcription and has been deployed for CRISPR activation (CRISPRa) (Hilton et al., 2015). The Krüppel-associated box (KRAB) domain is a transcriptional repressor that recruits other factors to methylate histones and has been used for CRISPR interference (CRISPRi) (Gilbert et al., 2013).

We began by interrogating the role of FACT in dCas9-based histone acetylation. We generated HEK293T cells that stably express both dCas9-p300 and a gRNA targeting the *CD25* TSS on either the coding or non-coding strand (Supplemental Figure 4A and Supplemental Table 2). Also known as *IL2RA*, *CD25* encodes a subunit of the interleukin-2 receptor and is poorly expressed under basal conditions (Uhlen et al., 2015), but its expression can be induced using CRISPRa (Simeonov et al., 2017). We transfected NT or SPT16 siRNAs into dCas9-p300 cells expressing either a coding or non-coding gRNA. Relative to the NT control, SPT16 depletion induced a significant increase in H3K27ac levels at the relevant site according to qPCR using primers (Supplemental Table 5) that amplified either a region upstream (Figure 4A) or inclusive (Supplemental Figure 4B) of the protospacer. Knockdown of SPT16 did not increase basal histone acetylation when dCas9-p300 was paired with a NT gRNA (Figure 4A and Supplemental Figure 4B).

**Figure 4.**
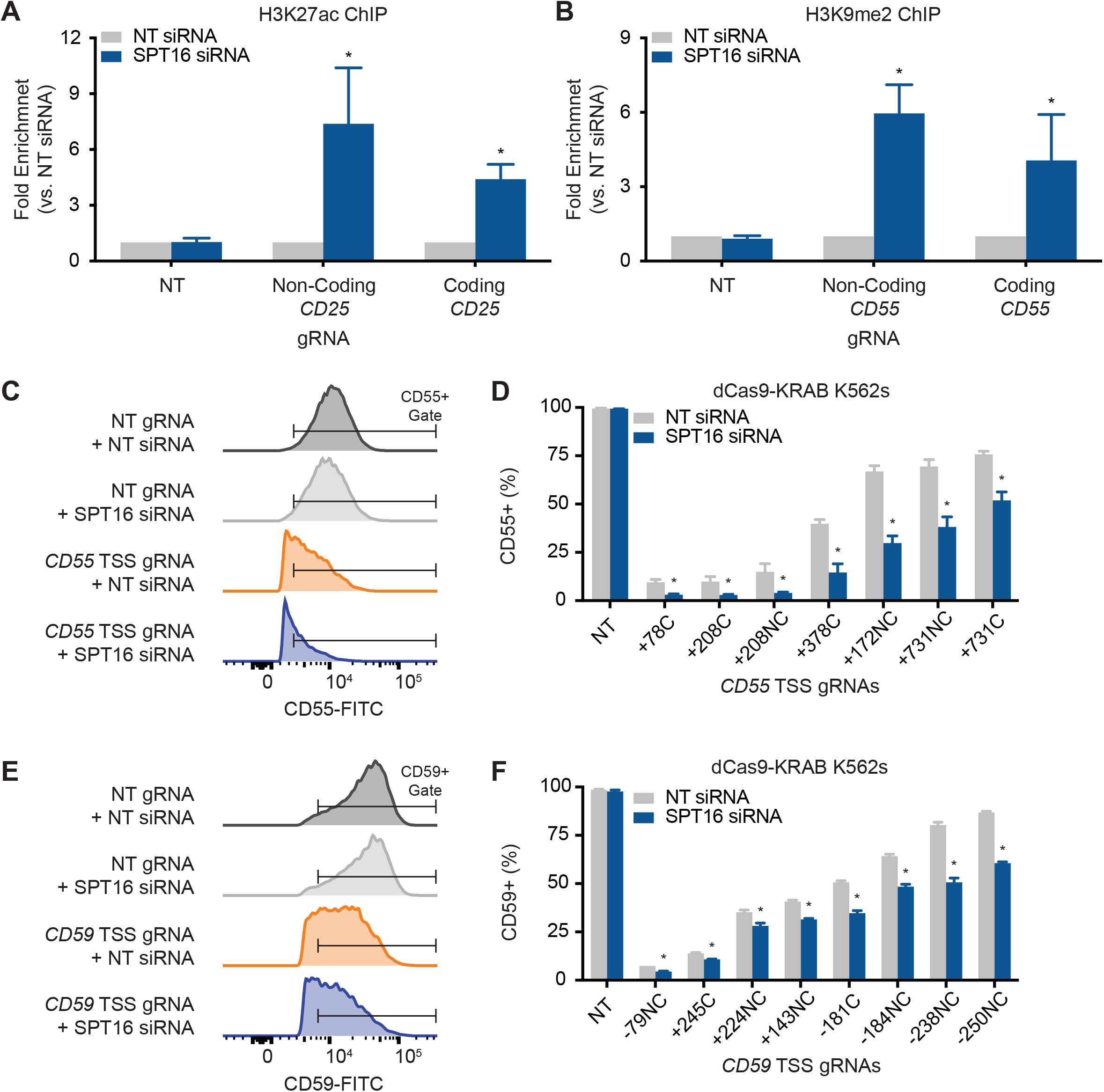
FACT Depletion Increases Epigenetic Marking and Transcriptional Phenotypes From dCas9-based Effectors in Human Cells. (A) Knockdown of SPT16 increases H3K27 acetylation in HEK29T dCas9-p300 cells (N = 3 biological replicates). Fold enrichment is the amount of H3K27ac after SPT16 depletion normalized to the amount of H3K27ac after treatment with a NT siRNA. qPCR primers amplified a region 9 base pairs upstream of the non-coding-strand gRNA protospacer and 46 base pairs upstream of the coding strand gRNA protospacer. (B) Knockdown of SPT16 increases H3K9 methylation in K562 dCas9-KRAB cells (N = 3 biological replicates). Fold enrichment is the amount of H3K9me2 after SPT16 depletion normalized to the amount of H3K9me2 after treatment with a NT siRNA. qPCR primers amplified a region 66 base pairs upstream of both the coding and non-coding strand gRNA protospacers. (C) Representative histograms of CD55 levels in CRISPRi cells after treatment with NT or SPT16 siRNAs. (D) FACT depletion enhances dCas9-KRAB-mediated knockdown of *CD55* in K562 cells expressing *CD55* TSS gRNAs (N = 3 biological replicates). CRISPRi cells were stained with CD55-FITC after transfection of either NT or SPT16 siRNAs. gRNAs bind to either the coding (C) or non-coding (NC) strand and are labeled according to their distance in base pairs from the TSS. (E) Representative histograms of CD59 levels in CRISPRi cells after treatment with NT or SPT16 siRNAs. (F) FACT depletion enhances dCas9-KRAB-mediated knockdown of *CD59* in K562 cells expressing *CD59* TSS gRNAs (N = 3 biological replicates). CRISPRi cells were stained with CD59-FITC after transfection of either NT or SPT16 siRNAs. gRNAs are labeled as in (D).

We used a similar approach to interrogate the role of FACT in dCas9-based histone methylation with K562 cells stably expressing dCas9-KRAB. We took advantage of conveniently located PAMs to target dCas9-KRAB to either the coding or non-coding strand of an identical location at the *CD55* TSS (Supplemental Figure 4C). *CD55* is a ubiquitously and highly expressed gene (Uhlen et al., 2015) that encodes a cell surface glycoprotein involved in the complement system. Transfection of SPT16 siRNAs increased levels of H3K9 methylation when dCas9-KRAB was targeted to either the coding or non-coding strand (Figure 4B, Supplemental Figure 4D, and Supplemental Table 5). Knockdown of SPT16 did not increase basal histone methylation when dCas9-KRAB was paired with a NT gRNA (Figure 4B and Supplemental Figure 4D).

We asked whether increased epigenetic marking by dCas9-based effectors during knockdown of FACT translates into increased transcriptional phenotypes. Targeting dCas9-p300 to the *CD25* TSS with gRNAs at varying distances from the TSS (Supplemental Table 2) generated cell populations that were between 12% and 95% CD25-positive, but knockdown of SPT16 did not further increase *CD25* expression for any gRNA tested (Supplemental Figure 5A and Supplemental Figure 6).

Targeting dCas9-KRAB to the *CD55* TSS with gRNAs at varying distances from the TSS (Supplemental Table 2) generated cell populations that were between 8% and 77% CD55-positive (Figures 4C, Figure 4D, and Supplemental Figure 7). Notably, siRNA knockdown of SPT16 potentiated the degree of CRISPRi as measured by a decrease in CD55-positive cells (Figures 4C and 4D) with some CRISPRi cell populations exhibiting more than a 60% drop in CD55-positive cells upon SPT16 depletion. This transcriptional phenotype was observed regardless of the strand to which the gRNA bound. We found similar potentiation of CRISPRi transcriptional phenotypes after SPT16 knockdown when targeting *CD59* (Figures 4E, Figure 4F, and Supplemental Figure 8), which is even more highly expressed on the cell surface than *CD55* (Uhlen et al., 2015). Increased CRISPRi was still dependent on proper targeting of dCas9-KRAB to the typical CRISPRi window around each gene’s TSS (Gilbert et al., 2013) as targeting dCas9-KRAB several kilobase pairs downstream of the TSS was ineffective even during knockdown of FACT (Supplemental Figure 5B). Similarly, localizing dCas9 unattached to an effector at various distances from the *CD55* or *CD59* TSSs did not affect transcription even after SPT16 knockdown (Supplemental Figures 5C-D, 9, and 10).

## Discussion

Programmable prokaryotic nucleases such as Cas9 are widely used for eukaryotic genome and transcriptome manipulation, but it is largely unclear how host cells interface with these foreign enzymes. While several studies have uncovered how histones impede Cas9 target search and binding (Hilton et al., 2015; Horlbeck et al., 2016; Kallimasioti-Pazi et al., 2018; Knight et al., 2015; Yarrington et al., 2018), none have described how proteins responsible for restructuring and remodeling nucleosomes might affect Cas9. We have found that the histone chaperone FACT is required for Cas9 unloading and multi-turnover activity in cell-free extract. In live human cells, knockdown of FACT inhibits templated repair of Cas9-induced breaks, increases indel formation, and increases the efficacy of dCas9-based transcriptional effectors. Our data do not rule out the potential importance of other histone chaperones or chromatin remodelers in genome surveillance and Cas9 displacement but highlight a prominent role of FACT in this process.

Modulating the turnover of Cas9 from a eukaryotic genome could inform the extent to which genome editing and transcriptional regulation rely upon repeated rounds of Cas9 binding and eviction. A kinetic analysis of Cas9 break repair suggested that cells primarily invoke error-prone pathways to slowly repair Cas9 breaks in a single round (Brinkman et al., 2018). Conversely, experiments inducing adjacent Cas9 breaks or modulating DNA repair with non-homologous single-stranded DNA implied that cells invoke error-free repair pathways that enable repeated rounds of Cas9 binding and eviction preceding eventual end-point mutation (Guo et al., 2018; Richardson et al., 2016a). We found that knockdown of FACT reduces Cas9-induced HDR and increases indels. NHEJ is a relatively fast process, relying on pathways that operate independent of the cell cycle. HDR alleles instead rely on slower pathways that depend upon passage through S/G2 (Hustedt and Durocher, 2016; Mao et al., 2008; Ranjha et al., 2018). Our data could imply that the slow appearance of HDR during Cas9 editing requires multiple rounds of Cas9 cleavage and turnover. In this scenario, knockdown of FACT could reduce the number of “shots on goal” available to the slower process of HDR, leaving cells to repair the break with a default error-prone pathway. However, we note that histone chaperones and chromatin remodelers can also directly influence DNA repair through multiple mechanisms such as increasing accessibility of the lesion to repair factors and promoting end-resection (Aleksandrov et al., 2018; Ayrapetov et al., 2014; Gospodinov et al., 2011; Lademann et al., 2017; Lans et al., 2012; Piquet et al., 2018; Price and D’Andrea, 2013). Hence, it is still unclear whether Cas9 owes its genome editing prowess to single-turnover or multi-turnover kinetics.

While Cas9 unloading within *X. laevis* egg extracts is ATP-dependent, FACT activity is ATP-independent (Orphanides et al., 1998). Notably, we found that depletion of ATP or immunodepletion of FACT are both sufficient to abrogate the multi-turnover behavior of Cas9 in *Xenopus* extract. It is possible that egg extracts actively recruit FACT to DNA-bound Cas9 and dCas9 in an ATP-dependent manner. We note that the most enriched factor in our proteomics data set is the lipid kinase PIP4K2C, and recent work has revealed that phosphoinositides localize around DNA lesions to recruit proteins through the lipids’ associations with Pleckstrin Homology (PH) domains (Wang et al., 2017). Intriguingly, both SPT16 and SSRP1 contain PH domains (Kemble et al., 2013; Zhang et al., 2015), but a great deal of additional work is necessary to determine if lipid metabolism around the R-loop generated by Cas9 binding is responsible for recruitment of FACT to Cas9.

While RNA polymerases are capable of dislodging Cas9 in a strand-specific manner *in vitro* (Clarke et al., 2018), our data argue that FACT plays a prominent role in Cas9 removal within eukaryotic systems. While FACT is often associated with transcription (Mason and Struhl, 2003; Saunders et al., 2003), it possesses nucleosome remodeling activity separate from RNA polymerase. The egg extract we employed is transcriptionally silent and does not initiate DNA replication (Lebofsky et al., 2009), indicating that basal FACT activity decoupled from transcription or replication within egg extract may be sufficient to remove Cas9 from its substrate. While previous studies have reported RNA polymerase-mediated displacement of Cas9 bound to the non-coding strand (Clarke et al., 2018), we and others have found no strand bias in dCas9-based epigenetic reprogramming (Gilbert et al., 2013; Hilton et al., 2015; Konermann et al., 2015; Qi et al., 2013).

Targeting dCas9 downstream of a TSS in *E. coli* effectively suppresses gene expression presumably because dCas9 acts as a potent transcriptional roadblock to the bacterial RNA polymerase (Qi et al., 2013). Transcriptional reprogramming in human cells is ineffective with dCas9 and requires recruitment of an epigenetic modifier (Gilbert et al., 2013; Qi et al., 2013), suggesting that dCas9 is not a roadblock to human RNA polymerases. We found that depleting FACT increased epigenetic marking and CRISPRi phenotypes for both coding and non-coding gRNAs across multiple loci. Notably, we also found that depleting FACT was insufficient to turn dCas9 alone into a transcriptional roadblock, further indicating that human RNA polymerases can displace Cas9. These results are consistent with Cas9’s utility as a generalized genome editing tool effective at editing both highly transcribed genes and transcriptionally silent regions.

FACT’s Cas9-displacing activity markedly influences epigenetic reprogramming by dCas9-fused effectors. Although FACT depletion did not increase CRISPRa transcriptional phenotypes in dCas9-p300 cells, knockdown of FACT induced an up to 7-fold increase in histone acetylation. Recruiting large numbers of transcriptional modulators successfully upregulates transcription (Chavez et al., 2015; Konermann et al., 2015; Tanenbaum et al., 2014), suggesting that dCas9-p300’s residence time may not be the critical bottleneck to increased expression. By contrast, increasing dCas9-KRAB’s residence time by globally downregulating FACT potentiates both dCas9-based histone methylation and transcriptional down-regulation. This down-regulation was evident only when we localized the KRAB domain around TSSs. Together with prior observations that Cas9-effectors are ineffective as short-lived RNPs but potent when expressed through permanent lentiviral constructs, our data suggest that the effectiveness of CRISPRi depends upon dCas9’s residence time at a TSS. Future approaches might specifically increase residence time without affecting other genome transactions.

## Acknowledgements

A.S.W. is supported by the National Science Foundation Graduate Research Fellowships Program and the Li Ka Shing Foundation. R.A.W. is supported by postdoctoral fellowship 131415-PF-17-168-01-DMC from the American Cancer Society. C.D.R., B.G.G., K.R.K., J.T.V., S.K.W., J.S., and J.E.C. are supported by from the Li Ka Shing Foundation and Heritage Medical Research Institute. J.T.V. is supported by CIRM TRAN1-09292. J.C.W. is supported by NIH grant HL098316. J.C.W. is a Howard Hughes Medical Institute Investigator and an American Cancer Society Research Professor. This work used the Vincent J. Coates Genomics Sequencing Laboratory at UC Berkeley, supported by NIH S10 OD018174 Instrumentation Grant. We would like to thank Dr. Danny Reinberg for generously providing recombinant FACT and Dr. Hasan Yardimci for generously providing the SPT16 antibody.

## Author Contributions

Conceptualization, A.S.W., R.A.W., C.D.R., J.C.W. and J.E.C.; Methodology, A.S.W., R.A.W., C.D.R., B.G.G., J.S., J.C.W. and J.E.C.; Software, S.K.W.; Formal Analysis, J.E.C.; Investigation, A.S.W., L.C., R.A.W., C.D.R., B.G.G., K.R.K., J.T.V., and S.K.W.; Writing – Original Draft, A.S.W. and J.E.C.; Writing – Review & Editing, A.S.W. and J.E.C.; Visualization, A.S.W.; Supervision, J.C.W. and J.E.C.; Funding Acquisition, J.C.W. and J.E.C.

## Declaration of Interests

J.E.C. is a co-founder of Spotlight Therapeutics.

---

**Figure S1 (Related to Figure 1): Purification and Activity of Recombinant Cas9-BirA* and dCas9-BirA\***

(A) Coomassie of purified Cas9, dCas9, Cas9-BirA*, and dCas9-BirA*.

(B) BirA* fusion does not compromise gRNA-targeting of Cas9 or Cas9’s cleavage ability *in vitro*.

(C) BirA* fusion does not impede energy-dependent removal of Cas9 from plasmid substrates in HSS. Incubation in buffer did not enable removal of dCas9-BirA* from DNA while incubation in HSS containing the ARS removed dCas9-BirA* from DNA to allow Cas9 cleavage. Pre-treating HSS with CIP and ATPγS inhibited dCas9-BirA* dislodging.

**Figure S2 (Related to Figure 2): Interactors of DNA-Bound Cas9 and dCas9**

(A) Proteins identified by mass spectrometry plotted according to average number of spectral counts in Cas9-BirA*–on-target gRNA samples (N = 3 biological replicates) versus dCas9-BirA*–on-target gRNA samples (N = 3 biological replicates).

(B) Immunodepletion of SSRP1 or SPT16 from HSS.

**Figure S3 (Related to Figure 3): Effects of FACT Depletion and ssODN Inclusion on Cas9 Editing Outcomes**

(A) Western blot of SPT16, SSRP1, and GAPDH in K562 cells transfected with either NT or SPT16 siRNAs.

(B) Indel rates from amplicon-NGS sequencing of eight different loci after electroporation of Cas9 RNPs in the absence of an HDR donor (N = 3 biological replicates).

(C) Total editing rates from amplicon-NGS sequencing of eight different loci after electroporation of Cas9 RNPs in the absence or presence of an HDR donor (N = 3 biological replicates).

(D) Total editing rates from amplicon-NGS sequencing of eight different loci after electroporation of Cas9 RNPs in the presence of an HDR donor (N = 3 biological replicates).

(E) Total editing rates from amplicon-NGS sequencing of *VEGFA* 0, 1, 2, 3, 6, 9, 12, and 24 hours after electroporation of Cas9 RNPs in the presence of an HDR donor (N = 3 biological replicates).

(F) Representative alleles from cells edited with Cas9 programmed with the *VEGFA* gRNA and a PAM-out ssODN HDR donor.

**Figure S4 (Related to Figure 4): FACT Depletion Enhances Chromatin Marking by dCas9-p300 and dCas9-KRAB**

(A) Schematic of coding and non-coding strand *CD25* gRNAs.

(B) Knockdown of SPT16 increases H3K27 acetylation in HEK29T dCas9-p300 cells (N = 3 biological replicates). Fold enrichment is the amount of H3K27ac after SPT16 depletion normalized to the amount of H3K27ac after treatment with a NT siRNA. qPCR primers amplified regions that include the corresponding protospacer.

(C) Schematic of coding and non-coding strand *CD55* gRNAs.

(D) Knockdown of SPT16 increases H3K9 methylation in K562 dCas9-KRAB cells (N = 3 biological replicates). Fold enrichment is the amount of H3K9me2 after SPT16 depletion normalized to the amount of H3K9me2 after treatment with a NT siRNA. qPCR primers amplified a region that include the protospacers.

**Figure S5 (Related to Figure 4): Enhancement of Transcriptional Engineering After FACT Depletion Requires Localizing KRAB domain to TSSs**

(A) FACT depletion does not affect *CD25* expression in HEK293T dCas9-p300 cells expressing *CD25* TSS gRNAs (N = 3 biological replicates). CRISPRa cells were stained with CD25-PE after transfection of either NT or SPT16 siRNAs. gRNAs are labeled as in Figure 4D.

(B) Knockdown of FACT does not affect *CD55* (left) or *CD59* (right) expression in K562 dCas9-KRAB cells expressing *CD55* or *CD59* gene body gRNAs (N = 3 biological replicates). gRNAs are labeled as in Figure 4D.

(C) Knockdown of FACT does not affect *CD55* expression in K562 dCas9 cells expressing *CD55* TSS gRNAs (N = 3 biological replicates). gRNAs are labeled as in Figure 4D.

(D) Knockdown of FACT does not affect *CD59* expression in K562 dCas9 cells expressing *CD59* TSS gRNAs (N = 3 biological replicates). gRNAs are labeled as in Figure 4D.

**Figure S6 (Related to Figure 4): Histograms of CD25 Levels in dCas9-p300 Cells After SPT16 Knockdown**

**Figure S7 (Related to Figure 4): Histograms of CD55 Levels in dCas9-KRAB Cells After SPT16 Knockdown**

**Figure S8 (Related to Figure 4): Histograms of CD59 Levels in dCas9-KRAB Cells After SPT16 Knockdown**

**Figure S9 (Related to Figure 4): Histograms of CD55 Levels in dCas9 Cells After SPT16 Knockdown**

**Figure S10 (Related to Figure 4): Histograms of CD59 Levels in dCas9 Cells After SPT16 Knockdown**

## STAR Methods

### Lead Contact and Materials Availability

Further information and requests for reagents and resources should be directed to the Lead Contact, Jacob Corn.

### *X. laevis* HSS

*X. laevis* HSS was prepared as previously described (Lebofsky et al., 2009). Aliquots were snap frozen and thawed as necessary. To immunodeplete SSRP1, HSS diluted 1:10 in Unloading Buffer (20 mM Tris pH 7.5, 100 mM KCl, 5 mM MgCl_2_, 1 mM DTT, 0.01% Tween) was exposed to two rounds of SSRP1 antibody. For each round, 150 μl of Dynabeads Protein G (Thermo Fisher Scientific) was washed three times in PBS, resuspended in 35 μg of SSRP1 antibody (Santa Cruz Biotechnology # sc-74536) and 825 μl of PBS, incubated with rotation for 90 min at room temperature, washed three times with PBST, resuspended in 2.4 μl of HSS diluted 1:10 in Unloading Buffer to 24 μl, and mixed for 45 min at room temperature. SPT16 antibody was generously provided by Dr. Hasan Yardimci. To immunodeplete SPT16, HSS was incubated with two rounds of 200 μl SPT16 antibody conjugated to 150 μl of Dynabeads Protein G. Mock depletions were conducted with the same amount of IgG antibody (Biolegend #400101).

### Cas9, FACT, RNA, and Donor DNA Preparation

Streptococcus pyogenes Cas9 (pMJ915, Addgene #69090) with two nuclear localization signal sequences and an HA tag at the C-terminus was expressed in Rosetta2 DE3 (UC Berkeley Macrolab) cells. Cell pellets were sonicated, clarified, Ni^2+^-affinity purified (HisTraps, GE Life Sciences), TEV cleaved, cation-exchanged (HiTrap SP HP, GE Life Sciences), size excluded (Sephacryl S-200, GE Life Sciences) and eluted at 40 μM in 20 mM HEPES KOH pH 7.5, 5% glycerol, 150 mM KCl, and 1 mM DTT. Recombinant human FACT was generously provided by Dr. Danny Reinberg. gRNAs were generated by HiScribe (New England Biolabs) T7 *in vitro* transcription using PCR-generated DNA as a template and purified using RNeasy Mini columns (Qiagen) (dx.doi.org/10.17504/protocols.io.dm749m). ssODN donor DNA was obtained by ordering unmodified ultramer oligonucleotides (Integrated DNA Technologies). For generation of stable cell lines, gRNAs were cloned into the lentiviral pGL1-library vector (Addgene #84832) as previously described (Horlbeck et al., 2016).

### Multi-Turnover Cas9 Activity

84 fmol of Cas9 diluted in Unloading Buffer to a volume of 1 μl was added to 504 fmol of gRNA diluted in Unloading Buffer to a volume of 0.5 μl. Cas9 and gRNA were incubated for 15 min at room temperature. 168 fmol of either linear or plasmid substrate was then added to the RNPs, and reaction mixtures were incubated for 45 min at room temperature. Next, either 3 μl of Unloading Buffer and 1 μl of 100 mM ATP, 3 μl of diluted HSS (1:8 in Unloading Buffer) and 1 μl of 100 mM ATP, 3 μl of mock-depleted and diluted HSS (1:8 in Unloading Buffer) and 1 μl of 100 mM ATP, 3 μl of SSRP1-depleted and diluted HSS (1:8 in Unloading Buffer) and 1 μl of 100 mM ATP, or 3 μl of diluted HSS (1:8 in Unloading Buffer) supplemented with CIP, ATPγS, and 1 μl of Unloading Buffer were added. 0.5 μl of CIP and 0.6 μl of ATPγS was added to every 18 μl of diluted HSS for the final condition. An additional 3 μl of Unloading Buffer and 1 μl of 100 mM ATP, 3 μl of diluted HSS (1:8 in Unloading Buffer) and 1 μl of 100 mM ATP, 3 μl of mock-depleted and diluted HSS (1:8 in Unloading Buffer) and 1 μl of 100 mM ATP, 3 μl of SSRP1-depleted and diluted HSS (1:8 in Unloading Buffer) and 1 μl of 100 mM ATP, or 3 μl of diluted HSS (1:8 in Unloading Buffer) supplemented with CIP and ATPγS and 1 μl of Unloading Buffer were added to the corresponding samples after 5, 10, 30, 60, 90, 120, and 150 min. Samples were mixed with Proteinase K (Sigma), incubated at 50°C for 30 min, and run on an agarose gel.

### Plasmid Protection in HSS

504 fmol of the on-target gRNA diluted in 0.5 μl of Unloading Buffer was added to either 0.5 μl of Unloading Buffer, 420 fmol of dCas9 diluted in 0.5 μl of Unloading Buffer, or 420 fmol of BirA*-dCas9 diluted in 0.5 μl of Unloading Buffer. Samples were incubated at room temperature for 15 min, added to 84 fmol of plasmid DNA diluted in 2.0 μl of Unloading Buffer, and incubated for 45 min at room temperature. Either 18 μl of Unloading Buffer, 18 μl of HSS supplemented with 0.55 μl of ARS, or 18 μl of HSS supplemented with 0.6 μl of ATPγS and 0.5 μl of CIP was added to the reaction mixtures. A stock solution of ARS was generated by mixing 10 μl of 100 mM ATP (VWR), 5 μl of 2 M phosphocreatine (Sigma-Aldrich), and 0.5 μl of 5 mg/ml creatine phosphokinase (Sigma-Aldrich). Samples were incubated for 15 min at room temperatures. Next, either 1 μl of Unloading Buffer or 4.2 pmol of Cas9 diluted in 0.5 μl of Unloading Buffer pre-complexed with 5.04 pmol of the on-target gRNA diluted in 0.5 μl of Unloading Buffer was added to the reaction mixtures and incubated for 30 min. Samples were incubated with Proteinase K (Sigma-Aldrich) at 50°C for 30 minutes and then run on an agarose gel. For the single-turnover protection assay using immunodepleted extracts, either 18 μl of diluted HSS (1:10 in Unloading Buffer) that was mock-depleted, 18 μl of diluted HSS (1:10 in Unloading Buffer) depleted of SSRP1, or 18 μl of diluted HSS (1:10 in Unloading Buffer) depleted of SPT16 was added to the RNPs after the 45 min incubation with the plasmid.

For FACT add-back experiments, 2 μg of recombinant human FACT was added to 18 μl of diluted HSS (1:10 in Unloading Buffer) immunodepleted of SSRP1 or SPT16 and incubated with pre-formed dCas9 RNP-plasmid complexes for 4 hours at room temperature prior to addition of Cas9 RNPs.

### Mass Spectrometry

5.04 pmol of the on-target gRNA diluted in 14 μl of Unloading Buffer was added to 4.2 pmol of either BirA*-Cas9 or BirA*-dCas9. As a reference sample, 5.04 pmol of a non-targeting gRNA diluted in 14 μl of Unloading Buffer was added to 4.2 pmol of BirA*-dCas9. RNPs were incubated at room temperature for 15 min. Samples were then added to 52.4 pmol of plasmid DNA, and reaction mixtures were incubated for 45 min at room temperature. 45 μl of HSS supplemented with 1.36 μl of ARS and biotin at a final concentration at 5 μM was added to these samples, and the resulting solutions were incubated for 60 min at room temperature. 405 μl of Unloading Buffer, 2.5 μl of Apyrase (New England Biolabs), and 56.25 μl of Apyrase Buffer was added to samples, which were incubated at 30°C for 15 minutes. 25 μl of DNase I (New England Biolabs) and 56.26 μl of DNase I Buffer were then added to samples, which were incubated at 37°C for 15 minutes. Samples were diluted 1:2 in Unloading Buffer and then mixed with 250 μl of MyOne™ Streptavidin C1 Dynabeads® (Thermo Fisher Scientific) that had been washed three times in 50 mM Tris, pH 7.4, 500 mM NaCl, 0.4% SDS, 5 mM EDTA, and 1 mM DTT. Samples were incubated overnight at 4°C with rotation.

Beads were washed once with 1 ml of 0.1% sodium deoxycholate, 1% Triton X-100, 500 mM NaCl, 1 mM EDTA, and 50 mM HEPES, pH 7.5, once with 1 ml of 250 mM LiCl, 0.5% NP-40, 0.5% sodium deoxycholate, 1 mM EDTA, 10 mM Tris, pH 8.1, and twice with 1 ml of 50 mM Tris, pH 7.4, 50 mM NaCl. Beads were then washed five times with 1 ml of 50 mM ammonium bicarbonate and then resuspended in 100 μl of 50 mM ammonium bicarbonate containing 0.01% ProteaseMAX (Promega) and 3 μg of sequencing-grade trypsin (Promega). Samples were incubated with mixing at 37°C for 4 hours after which the supernatant was collected and transferred to a new tube. Beads were washed again with 50 μl of 50 mM ammonium bicarbonate, and supernatants were pooled. 2 μl of formic acid (Fisher Scientific) was added to acidify the samples to a pH of ∼3.0. Samples were then spun down to dryness in a speedvac and submitted to the University of California, Davis Proteomics Core for Multi-Dimension Protein Identification Technology mass spectrometry. Trypsinized peptides were mapped to the *X. laevis* proteome using the PHROG database (Wuhr et al., 2014). Protein enrichment levels were analyzed by the Limma Bioconductor package.

### Western Blots

For *X. laevis* HSS, samples of equal volumes were incubated with Laemmli Buffer (Bio-Rad) at 95°C for 5 min. For human tissue culture, cells were washed in PBS and then lysed in 1X RIPA Buffer (Millipore Sigma) supplemented with Halt™ Protease Inhibitors (Thermo Fisher Scientific) at 4°C for 60 min. Samples were spun down at 15,000 g for 15 min, and the protein concentrations of the cleared lysates were measured using a BCA Protein Assay Kit (Thermo Fisher Scientific). 30 ug of lysate was denatured by incubation with Laemmli Buffer at 95°C for 5 min.

Both *X. laevis* and mammalian protein samples were resolved on Mini-PROTEAN^®^ TGX™ 4-20% gels (Bio-Rad), and resolved proteins were transferred (TransBlot Turbo, Bio-Rad) to nitrocellulose membranes. Membranes were blocked in 5% milk in TBST for 30 min at room temperature and incubated with primary antibodies in blocking buffer overnight at 4°C. Membranes were washed three times in TBST, incubated with secondary antibodies (LI-COR Biosciences) in blocking buffer for 45 min, and then exposed on an Odyssey^®^ CLx Imaging System (LI-COR Biosciences). *X. laevis* protein levels were probed using the following antibodies: GAPDH (Cell Signaling #2118 1:5000), SPT16 (Cell Signaling #12191 1:1000), and SSRP1 (Santa Cruz Biotechnology sc-74536 1:1000). Human cell protein levels were probed using the following antibodies: GAPDH (Cell Signaling #2118 1:5000), SPT16 (Cell Signaling #12191 1:1000), SSRP1 (Biolegend #609701 1:1000), and INO80B (Santa Cruz Biotechnology 1:1000).

### Cell Culture

HEK293T dCas9-p300 cells were a generous gift from Dr. Charles Gersbach. Parental K562 cells were acquired from the UC Berkeley Cell Culture Facility. K562 dCas9-KRAB were identical to those previously reported (Richardson et al., 2018). All cells were regularly tested for mycoplasma contamination. HEK293T cells were maintained in DMEM with glutamax (Gibco) supplemented with 10% fetal bovine serum, 1% sodium pyruvate (Gibco), and 100 U/ml penicillin-streptomycin (Gibco). K562 cells were maintained in RPMI (Gibco) supplemented with 10% fetal bovine serum, 1% sodium pyruvate, and 100 U/ml penicillin-streptomycin.

### Lentiviral Packaging and Transduction

Lentiviral packaging of all constructs was performed in HEK293T cells. Plasmids were transfected using TransIT®-LT1 Transfection Reagent (Mirus) at a ratio of 1 μg of total DNA to 3 μl of the transfection reagent. The plasmid mixture consisted of 50% lentiviral transfer plasmid, 40% ΔVPR plasmid, and 10% VSVG plasmid. Virus was harvested at 48 and 72 hours after transfection, passed through a 0.45 μM filter, and added to target cells for transduction. 48 hours after transduction, both K562 and HEK293T cells were exposed to puromycin at 1 μg/ml. Cells were maintained in media containing puromycin for at least two passages to ensure complete selection.

### siRNA Transfection

For Western blots and flow cytometry, 50,000 cells were transfected with 7.5 pmol of siRNA complexed with 2.25 μl of Lipofectamine™ RNAiMAX (Thermo Fisher Scientific) in Opti-MEM (Gibco). For ChIP experiments, 5,000,000 HEK293T or K562 cells were transfected with 750 pmol of siRNA complexed with 225 μl of Lipofectamine™ RNAiMAX in Opti-MEM. For editing experiments, 1,200,000 K562 cells were transfected 180 pmol of siRNA complexed with 54 μl of Lipofectamine™ RNAiMAX. Cells were transfected in the absence of penicillin-streptomycin. 12 hours after transfection, cells were transferred to fresh media containing penicillin-streptomycin. The following siRNAs were used: SMARTpool ON-TARGETplus SUPT16H siRNA (GE Dharmacon), SMARTpool ON-TARGETplus INO80B siRNA (GE Dharmacon), and ON-TARGETplus Nontargeting Pool (GE Dharmacon).

### Flow Cytometry

Sixty hours after transfection of siRNAs, cells were washed once in 1% BSA in PBS and then stained on ice for 1 hour. Cells were stained in 50 μl of either a PE CD25 antibody (Bioegend #302606 1:100), FITC CD55 antibody (Biolegend #311305 1:100), or FITC CD59 antibody Biolegend #304706). Samples were also stained with FITC Mouse IgG1, κ antibody (Biolegend #400107 1:100) as an isotype control. Cells were washed three times in 1% BSA in PBS. Fluorescence was measured using the Attune NxT Flow Cytometer (Thermo Fisher Scientific).

### Chromatin Immunoprecipitation

Sixty hours after transfections, cells were trypsinized if necessary, washed once in PBS, and then incubated in 10 ml of 1% formaldehyde in PBS for 15 min at room temperature. Reactions were quenched with 1.54 ml of 1.5 M glycine. Samples were spun down, resuspended in 1 ml of ice-cold PBS, spun down again, and snap-frozen in liquid nitrogen.

Cell pellets were lysed after thawing on ice by incubation first in 10 ml of 50 mM HEPES, 140 mM NaCl, 1 mM EDTA, 10% glycerol, 0.5% NP-40, 0.25% Triton X-100, and Halt™ Protease Inhibitors at 4°C for 10 min. Samples were spun down, the supernatant was aspirated, and the pellet was resuspended in 10 ml of 200 mM NaCl, 1 mM EDTA, 0.5 mM EGTA, 10 mM Tris, and Halt™ Protease Inhibitors at 4°C for 10 min. Samples were spun down again, the supernatant was aspirated, and the pellets were resuspended in 900 μl of 10 mM Tris, 1 mM EDTA, 0.5 mM EGTA, 100 mM NaCl, 0.1% sodium deoxycholate, 0.5% sarcosine, and Halt™ Protease Inhibitors. Resuspended samples were sonicated with a Sartorius probe sonicator with three 1-minute intervals with an amplitude of 70% and a cycle of 0.9. Sonicated samples were added to 5.1 ml of 10 mM Tris, 1 mM EDTA, 0.5 mM EGTA, 100 mM NaCl, 0.1% sodium deoxycholate, 0.5% sarcosine, Halt™ Protease Inhibitors, and 600 μl of 10% Triton X-100. Solutions were centrifuged at 4°C at max speed for 20 minutes. 150 μl of the supernatant was retained as input. 6 ml of supernatant was used for immunoprecipitation.

8 μg of H3K9me2 (Abcam ab1220), H3K27ac (Abcam ab4729), or IgG antibody (Biolegend 400101) was incubated with 100 μl of Protein G Dynabeads (Thermo Fisher Scientific) at room temperature for 2 hours, washed three times with 0.5% BSA in PBS, resuspended in 100 μl of 0.5% BSA in PBS, and added to 3 ml of cell lysate. Samples were incubated overnight at 4°C with rotation. Beads were then washed four times with 1 ml of 10 mM HEPES, 500 mM LiCl, 1 mM EDTA, 1% NP-40, and 0.7% sodium deoxycholate, washed once in 1 ml of TBS, resuspended in 200 μl of 50 mM Tris, 10 mM EDTA, 1% SDS, and incubated at 65°C overnight. Supernatants were collected and added to 200 μl of TE. 100 μl of input lysate was added to 300 μl of TE. 1 μl of RNAse A (New England Biolabs) was added to the samples, which were incubated at 37°C for 1 hour. 4 μl of Proteinase K (Thermo Fisher Scientific) was added to the samples, which were incubated at 57°C for 1 hour. 2 ml of Buffer PB (Qiagen) was added to the samples, which were flowed over MinElute columns (Qiagen). Columns were washed with Buffer PE (Qiagen) and eluted in 30 μl of Buffer EB (Qiagen). Immunoprecipitation samples were diluted 1:3 in distilled water while input samples were diluted 1:49 in distilled water.

qPCR reactions were performed in a total volume of 10 μl containing 2 μl of diluted samples, 5 ul of 2X ssoFast™ EvaGreen Supermix with Low ROX (Bio-Rad), and primers (Supplemental Table 5) each at a final concentration of 500 nM. Samples were run on an Applied Biosystems™ StepOne™ Real-Time PCR System (Fisher Scientific). The thermocycler was set for 95°C for 2 minutes and 40 cycles of 95°C for 5 seconds and 55°C for 10 seconds. Fold enrichment of the assayed genes over a control locus were calculated using the 2^−ΔΔCt^ method.

### Cas9 Electroporation

Cells were electroporated with Cas9 RNPs 60 hours after transfection of siRNAs. For each electroporation, 30 pmol of Cas9 was diluted to a final volume of 3.5 μl with Cas9 Buffer (20 mM HEPES pH 7.5, 150 mM KCl, 1 mM MgCl2 10% glycerol and 1 mM TCEP). Cas9 was incubated with 36 pmol of gRNA diluted to a final volume of 3.5 μl in Cas9 Buffer. The resulting mixture was incubated for 10 min at room temperature. For HDR experiments, 1 μl of 100 μM ssODN donor was then added to the RNPs. 200,000 cells were washed once in PBS, resuspended in 16 μl of Buffer SF (Lonza), added to the RNP complexes, and electroporated using the FF120 program on the 4D-Nucleofector™ (Lonza). Reaction mixtures were incubated at room temperature for 10 min after electroporation and then transferred to pre-warmed media.

### Next-Generation Sequencing

Either 0, 1, 2, 3, 6, 9, 12, 24 or 48 hours after electroporation, genomic DNA was harvested cells using QuickExtract™ DNA Extraction Solution (Lucigen). 200 ng of genomic DNA from edited cells was amplified using primer pairs from primer set 1 in a 30-cycle PCR reaction (Supplemental Table 4). PCR products were SPRI cleaned, and 25 ng of SPRI-cleaned amplicons were amplified again using primer pairs from primer set 2 (Supplemental Table 4) in a 12-cycle PCR reaction. Amplicons from the second PCR were SPRI cleaned, and 10 ng of SPRI-cleaned amplicons were used in a 9-cycle PCR reaction with Illumina compatible primers. All PCRs were conducted with PrimeSTAR GXL DNA Polymerase (Takara) according to the manufacturer’s instructions. Libraries were pooled and submitted to the Vincent J. Coates Genomics Sequencing Laboratory at the University of California, Berkeley for 300 bp paired-end cycle processing using an Illumina MiSeq sequencing Kit (Illumina Inc., San Diego, CA).

Samples were deep sequenced to a depth of at least 10,000 reads. Reads were trimmed of adapters and low quality bases. Paired reads were joined into a single read and aligned to the input reference and donor sequences using NEEDLE (Li et al., 2015). Editing outcomes were determined and quantified using a modified version (https://github.com/staciawyman/cortado) of CRISPResso (Pinello et al., 2016). Reads were classified as NHEJ if an insertion or deletion in the alignment overlapped a 6 bp window around the cut site. Reads were classified as HDR if they were not NHEJ and contained the primary edit specified in the donor sequence. Percent NHEJ and HDR are calculated as the number of reads divided by the number of aligned reads.

### Statistical Analysis

All analysis was performed using data from three biological replicates. Data are presented as mean ± standard deviation. Statistical analyses were performed in the PRISM software using a Student’s t-test.

