## Supplemental Figures and Text for "The histone chaperone FACT induces Cas9 multi-turnover behavior and modifies genome manipulation in human cells"

### Supplemental Figure 1

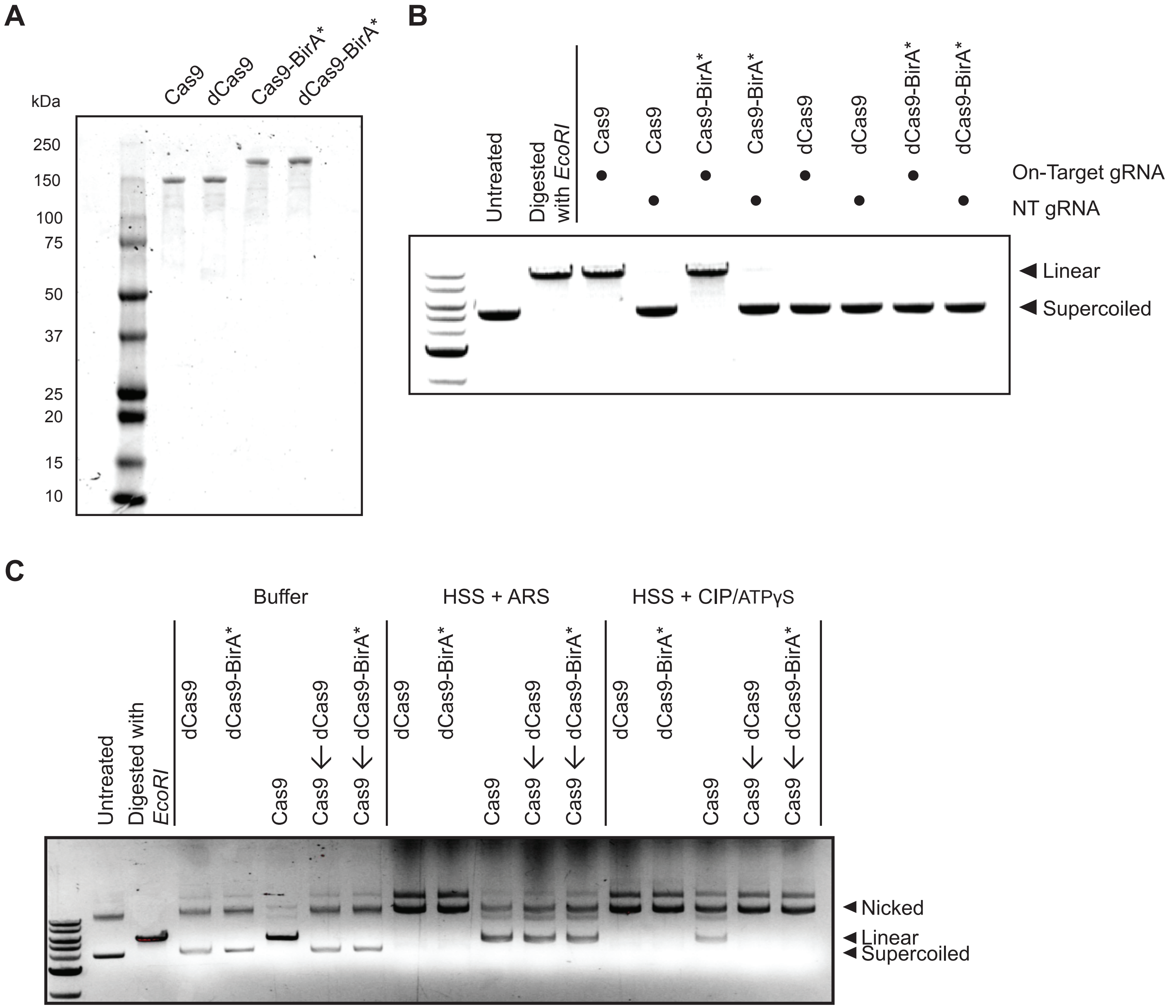

### Supplemental Figure 2

A

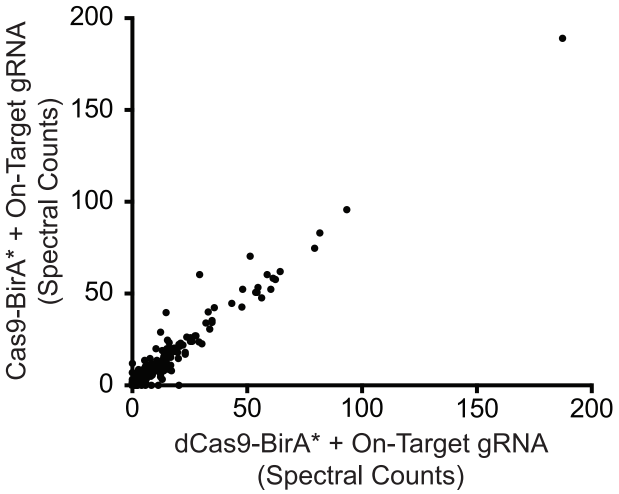


B

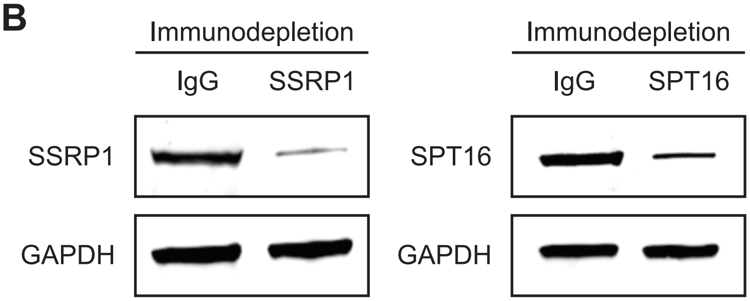

### Supplemental Figure 3

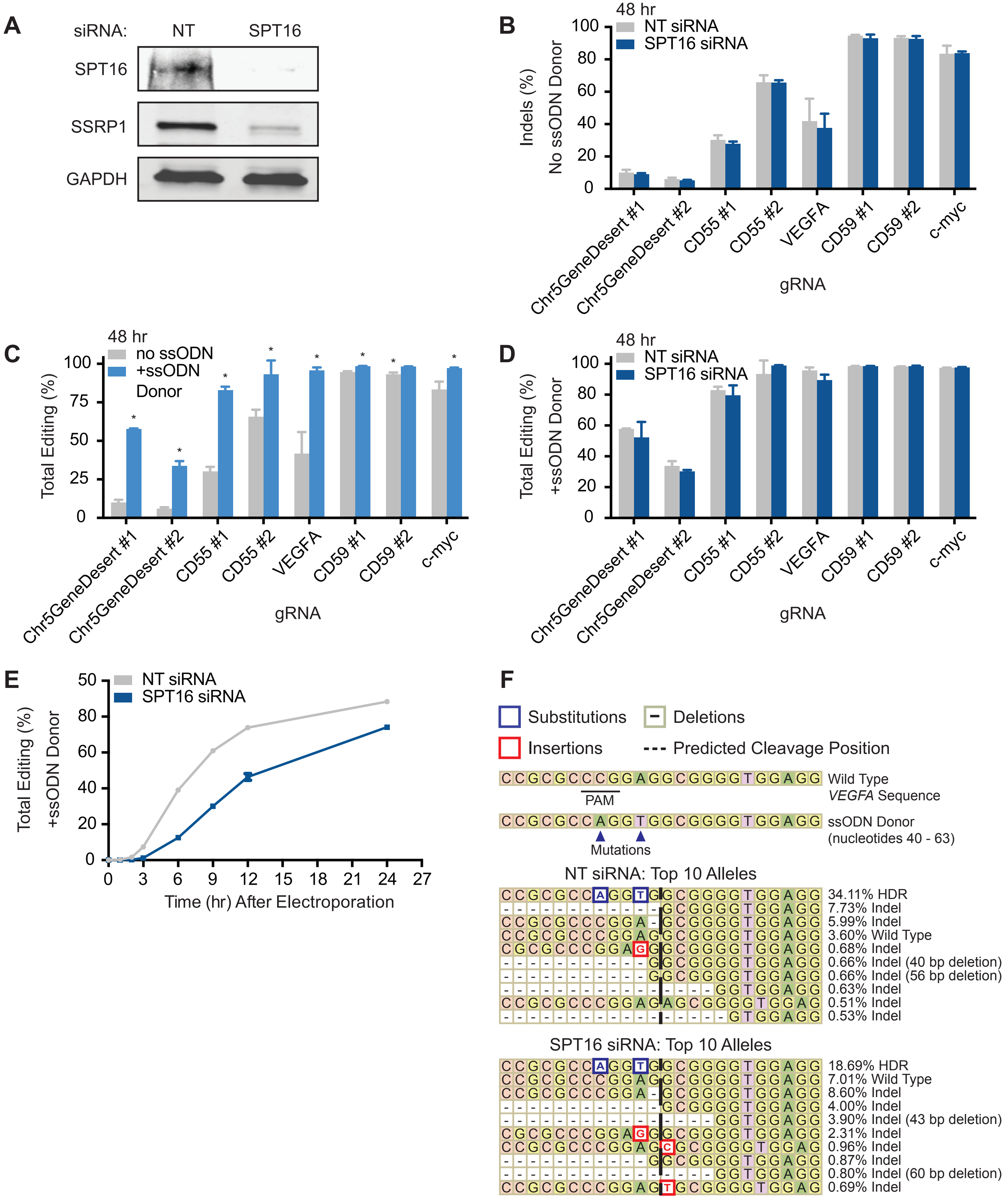

### Supplemental Figure 4

**A**

Coding Strand
*CD25* gRNA

5′ – CCTGGCTGAACACGCCAGCCCAA – 46 bp – CACAAGGGTGACAGCCCAGGCGG – 3′
3′ – GGACCGACTTGTGCGGTCGGGTT – 46 bp – GTGTTCCCACTGTCGGGTCCGCC – 5′

Non-Coding Strand
*CD25* gRNA

**B**

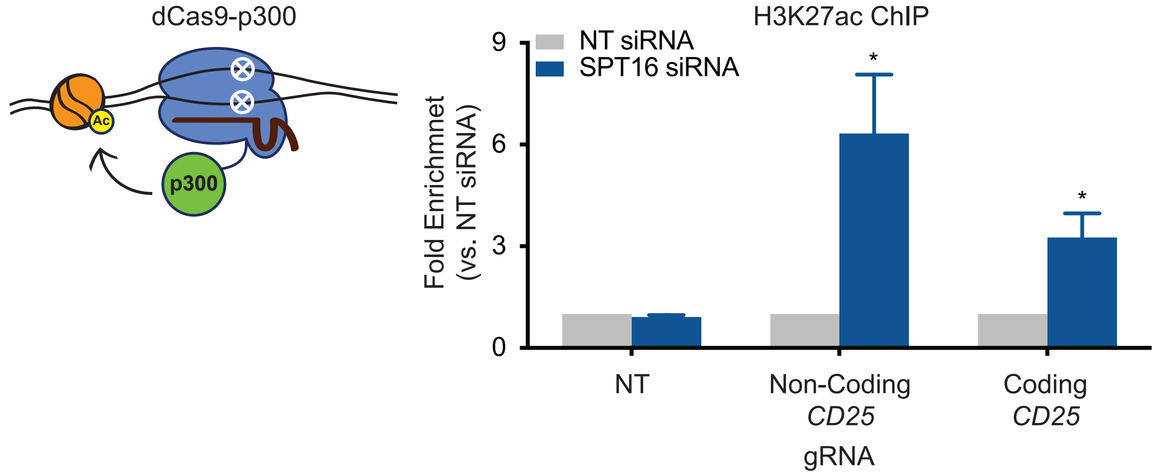


**C**

Coding Strand
*CD55* gRNA

5′ – CCTAATGCCCAGCCAGCTTTGGATGG – 3′
3′ – GGATTACGGGTCGGTCGAAACCTACC – 5′

Non-Coding Strand
*CD55* gRNA

**D**

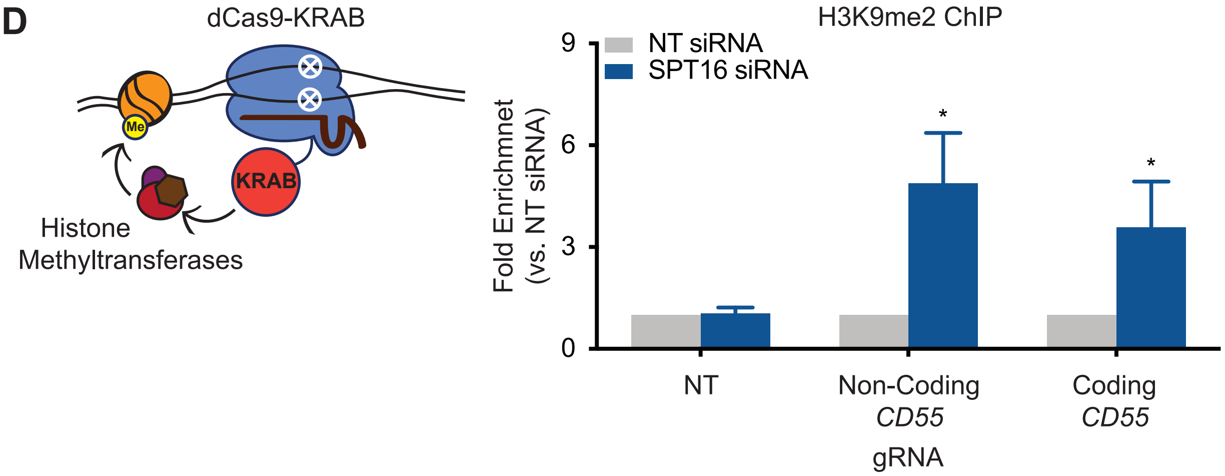

### Supplemental Figure 5

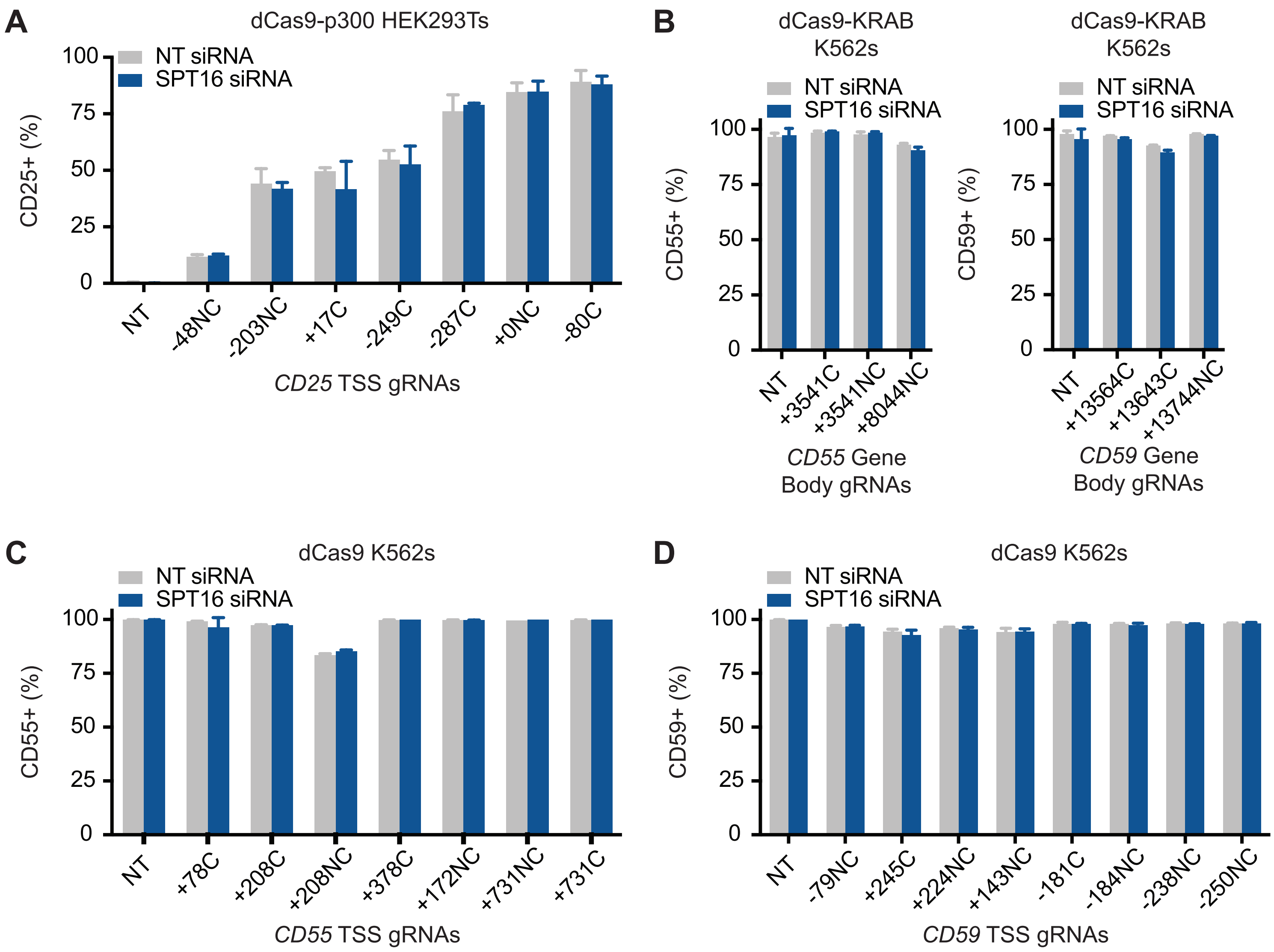

### Supplemental Figure 6

dCas9-p300 HEK293Ts - CD25 Levels

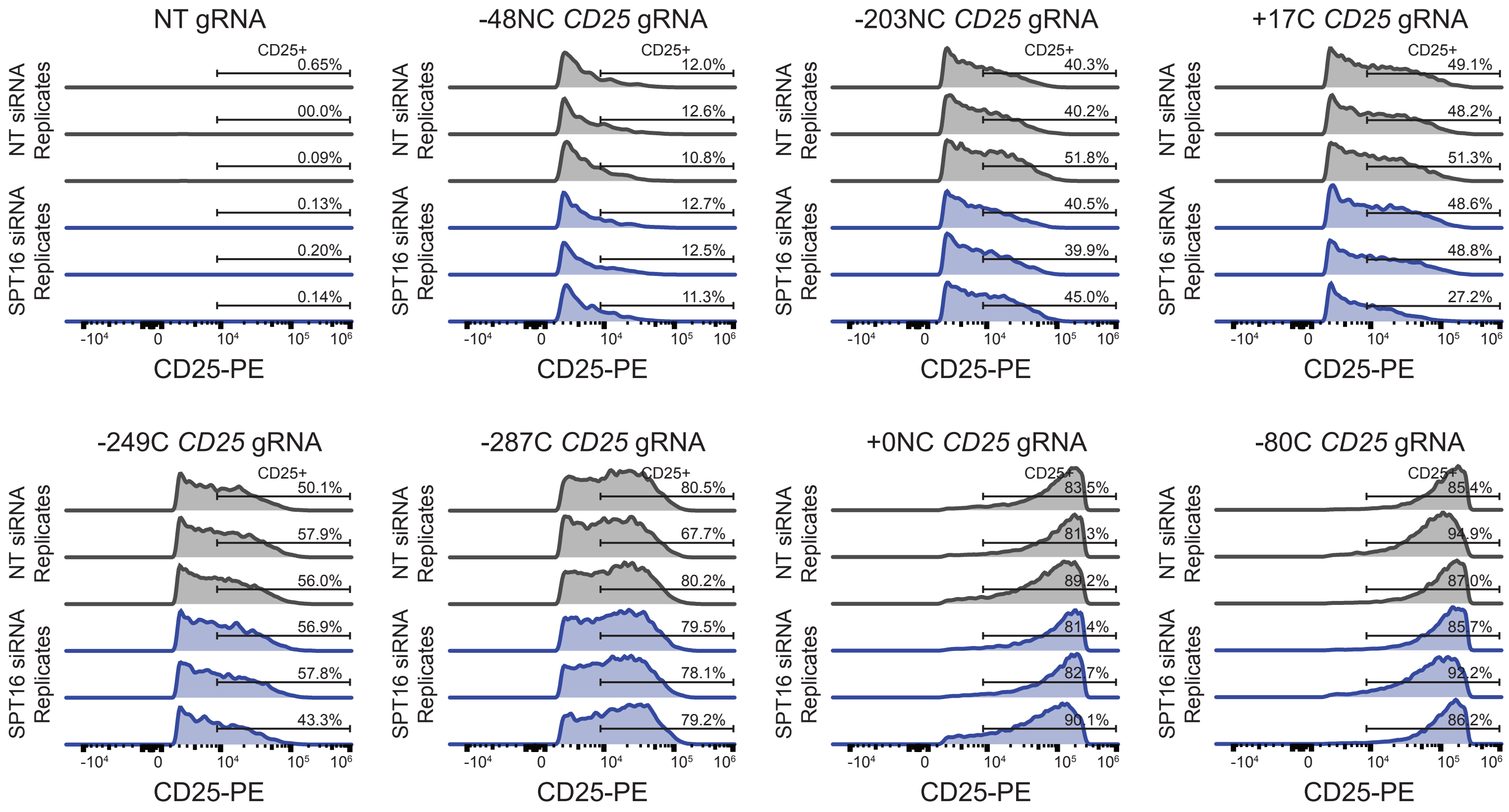

### Supplemental Figure 7

dCas9-KRAB K562s - CD55 Levels

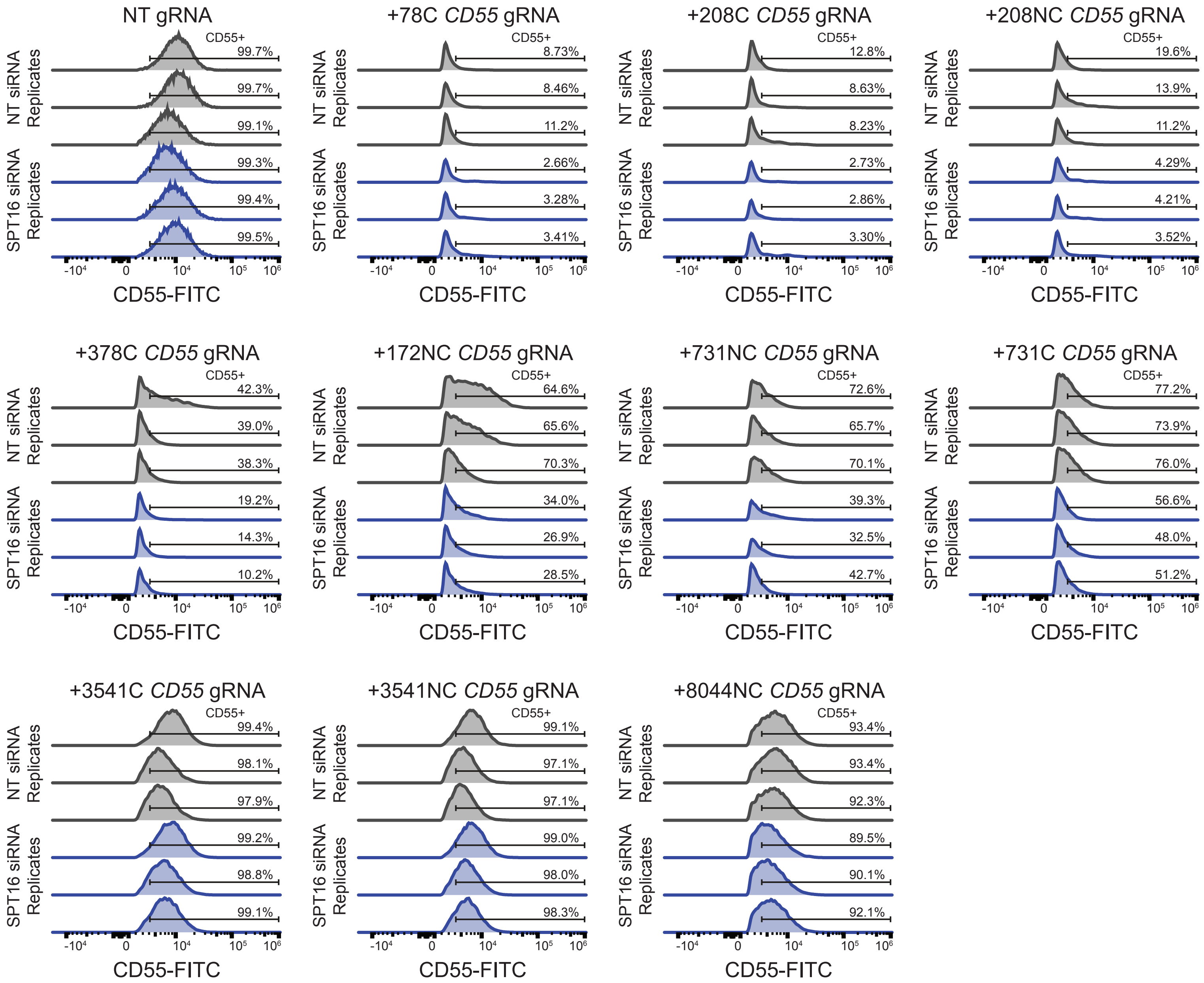

### Supplemental Figure 8

dCas9-KRAB K562s - CD59 Levels

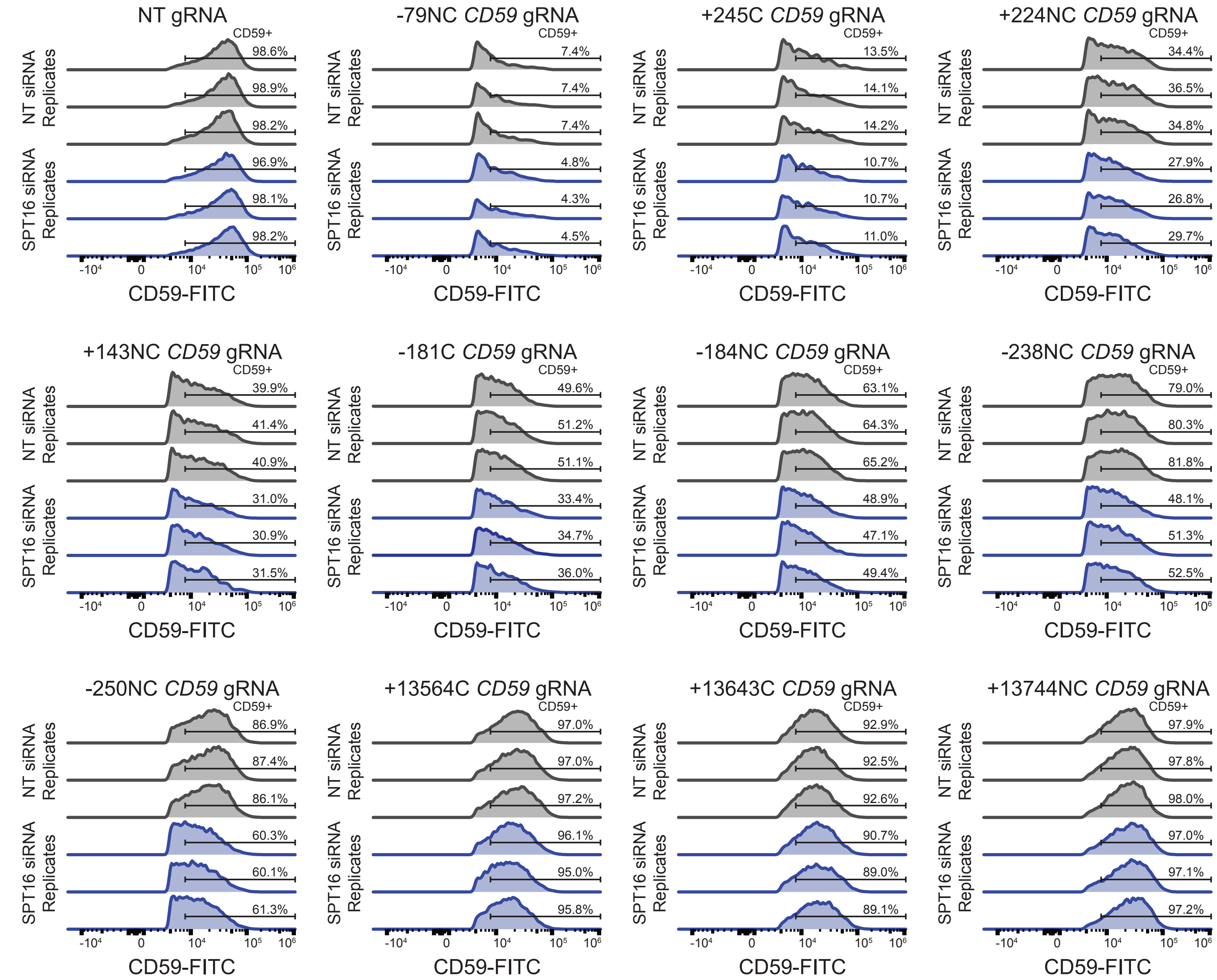

### Supplemental Figure 9

dCas9 K562s - CD55 Levels

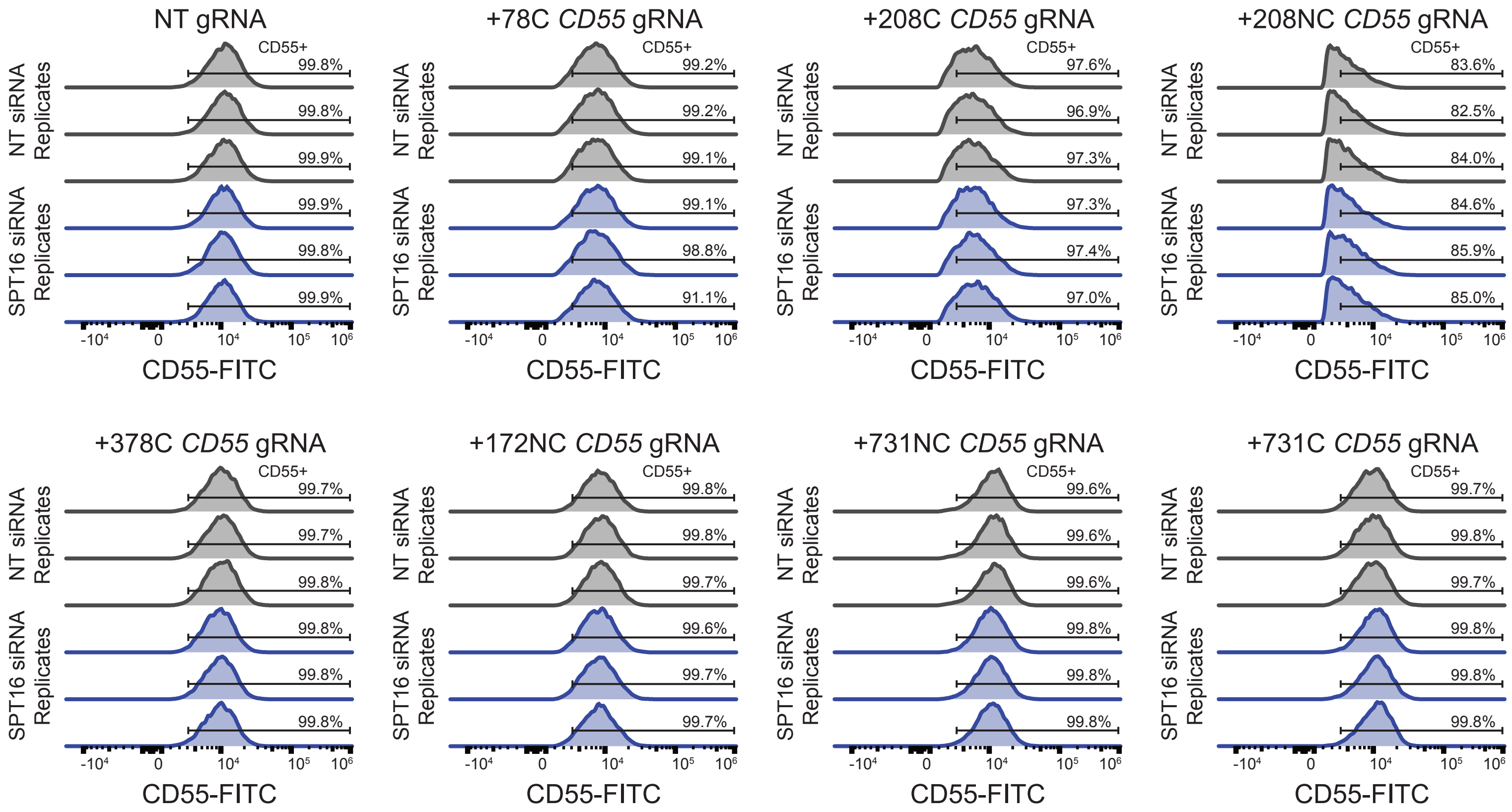

### Supplemental Figure 10

dCas9 K562s - CD59 Levels

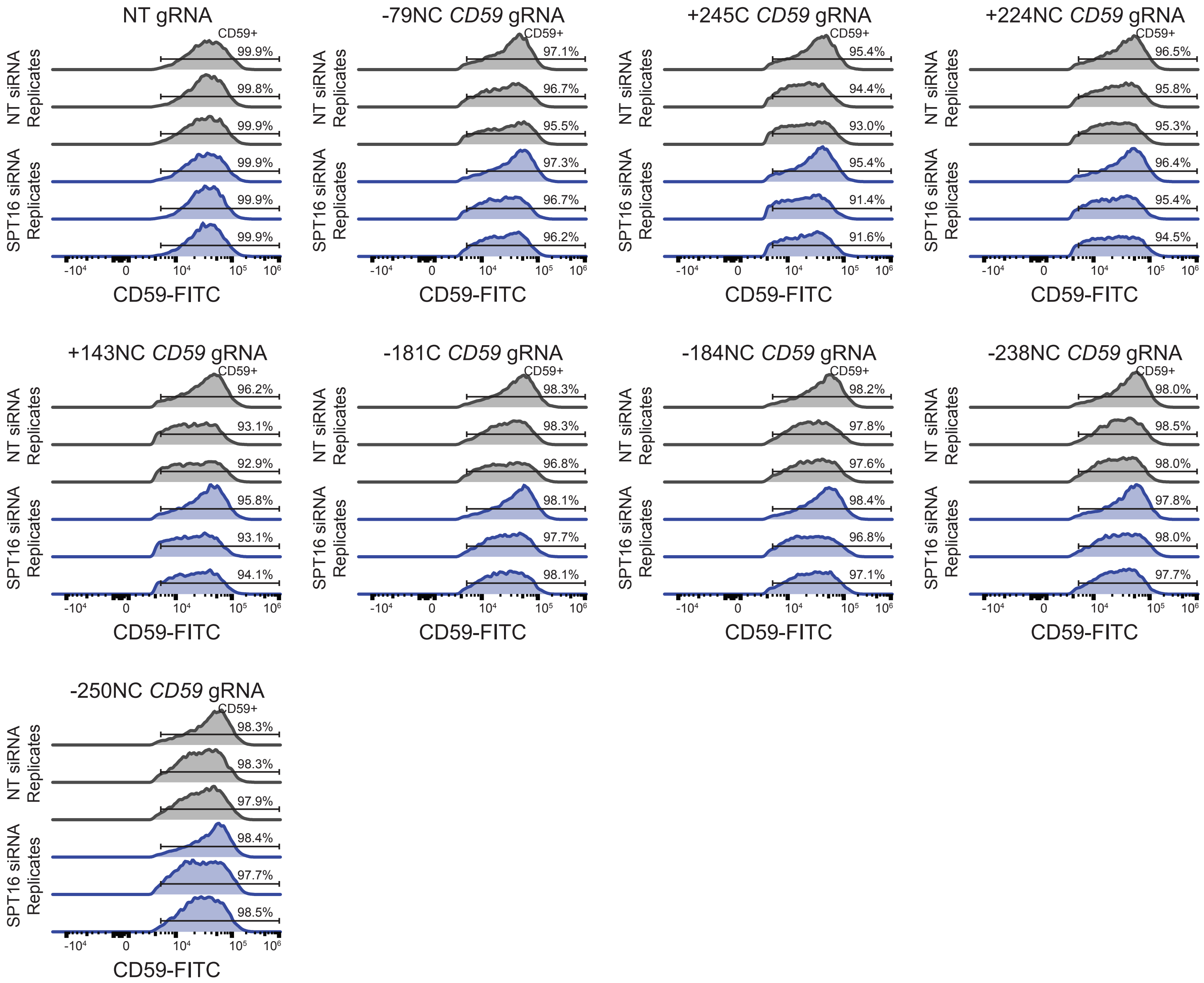

**Supplemental Information**

**Supplemental Data S1 (Related to Figure 2): Amino Acid Sequences of Cas9-BirA* and dCas9-BirA***

Cas9-BirA*:

MDYKDHDGDYKDHDIDYKDDDDKMAPKKKRKVGIHGVPAADKKYSIGLDIGTNSVGWAVITDE
YKVPSKKFKVLGNTDRHSIKKNLIGALLFDSGETAEATRLKRTARRRYTRRKNRICYLQEIFSNE
MAKVDDSFFHRLEESFLVEEDKKHERHPIFGNIVDEVAYHEKYPTIYHLRKKLVDSTDKADLRLI
YLALAHMIKFRGHFLIEGDLNPDNSDVDKLFIQLVQTYNQLFEENPINASGVDAKAILSARLSKSR
RLENLIAQLPGEKKNGLFGNLIALSLGLTPNFKSNFDLAEDAKLQLSKDTYDDDLDNLLAQIGDQ
YADLFLAAKNLSDAILLSDILRVNTEITKAPLSASMIKRYDEHHQDLTLLKALVRQQLPEKYKEIFF
DQSKNGYAGYIDGGASQEEFYKFIKPILEKMDGTEELLVKLNREDLLRKQRTFDNGSIPHQIHLG
ELHAILRRQEDFYPFLKDNREKIEKILTFRIPYYVGPLARGNSRFAWMTRKSEETITPWNFEEVV
DKGASAQSFIERMTNFDKNLPNEKVLPKHSLLYEYFTVYNELTKVKYVTEGMRKPAFLSGEQK
KAIVDLLFKTNRKVTVKQLKEDYFKKIECFDSVEISGVEDRFNASLGTYHDLLKIIKDKDFLDNEE
NEDILEDIVLTLTLFEDREMIEERLKTYAHLFDDKVMKQLKRRRYTGWGRLSRKLINGIRDKQSG
KTILDFLKSDGFANRNFMQLIHDDSLTFKEDIQKAQVSGQGDSLHEHIANLAGSPAIKKGILQTVK
VVDELVKVMGRHKPENIVIEMARENQTTQKGQKNSRERMKRIEEGIKELGSQILKEHPVENTQL
QNEKLYLYYLQNGRDMYVDQELDINRLSDYDVDHIVPQSFLKDDSIDNKVLTRSDKNRGKSDN
VPSEEVVKKMKNYWRQLLNAKLITQRKFDNLTKAERGGLSELDKAGFIKRQLVETRQITKHVAQ
ILDSRMNTKYDENDKLIREVKVITLKSKLVSDFRKDFQFYKVREINNYHHAHDAYLNAVVGTALIK
KYPKLESEFVYGDYKVYDVRKMIAKSEQEIGKATAKYFFYSNIMNFFKTEITLANGEIRKRPLIET
NGETGEIVWDKGRDFATVRKVLSMPQVNIVKKTEVQTGGFSKESILPKRNSDKLIARKKDWDP
KKYGGFDSPTVAYSVLVVAKVEKGKSKKLKSVKELLGITIMERSSFEKNPIDFLEAKGYKEVKKD
LIIKLPKYSLFELENGRKRMLASAGELQKGNELALPSKYVNFLYLASHYEKLKGSPEDNEQKQLF
VEQHKHYLDEIIEQISEFSKRVILADANLDKVLSAYNKHRDKPIREQAENIIHLFTLTNLGAPAAFK
YFDTTIDRKRYTSTKEVLDATLIHQSITGLYETRIDLSQLGGDKRPAATKKAGQAKKKKGGGSG
GGSGGGSKDNTVPLKLIALLANGEFHSGEQLGETLGMSRAAINKHIQTLRDWGVDVFTVPGKG
YSLPEPIQLLNAKQILGQLDGGSVAVLPVIDSTNQYLLDRIGELKSGDACIAEYQQAGRGGRGR
KWFSPFGANLYLSMFWRLEQGPAAAIGLSLVIGIVMAEVLRKLGADKVRVKWPNDLYLQDRKL
AGILVELTGKTGDAAQIVIGAGINMAMRRVEESVVNQGWITLQEAGINLDRNTLAAMLIRELRAA
LELFEQEGLAPYLSRWEKLDNFINRPVKLIIGDKEIFGISRGIDKQGALLLEQDGIIKPWMGGEISL
RSAEKAYPYDVPDYA*

dCas9-BIrA*:

MDYKDHDGDYKDHDIDYKDDDDKMAPKKKRKVGIHGVPAADKKYSIGLAIGTNSVGWAVITDE
YKVPSKKFKVLGNTDRHSIKKNLIGALLFDSGETAEATRLKRTARRRYTRRKNRICYLQEIFSNE
MAKVDDSFFHRLEESFLVEEDKKHERHPIFGNIVDEVAYHEKYPTIYHLRKKLVDSTDKADLRLI
YLALAHMIKFRGHFLIEGDLNPDNSDVDKLFIQLVQTYNQLFEENPINASGVDAKAILSARLSKSR
RLENLIAQLPGEKKNGLFGNLIALSLGLTPNFKSNFDLAEDAKLQLSKDTYDDDLDNLLAQIGDQ
YADLFLAAKNLSDAILLSDILRVNTEITKAPLSASMIKRYDEHHQDLTLLKALVRQQLPEKYKEIFF
DQSKNGYAGYIDGGASQEEFYKFIKPILEKMDGTEELLVKLNREDLLRKQRTFDNGSIPHQIHLG
ELHAILRRQEDFYPFLKDNREKIEKILTFRIPYYVGPLARGNSRFAWMTRKSEETITPWNFEEVV
DKGASAQSFIERMTNFDKNLPNEKVLPKHSLLYEYFTVYNELTKVKYVTEGMRKPAFLSGEQK
KAIVDLLFKTNRKVTVKQLKEDYFKKIECFDSVEISGVEDRFNASLGTYHDLLKIIKDKDFLDNEE
NEDILEDIVLTLTLFEDREMIEERLKTYAHLFDDKVMKQLKRRRYTGWGRLSRKLINGIRDKQSG
KTILDFLKSDGFANRNFMQLIHDDSLTFKEDIQKAQVSGQGDSLHEHIANLAGSPAIKKGILQTVK

VVDELVKVMGRHKPENIVIEMARENQTTQKGQKNSRERMKRIEEGIKELGSQILKEHPVENTQL
QNEKLYLYYLQNGRDMYVDQELDINRLSDYDVDAIVPQSFLKDDSIDNKVLTRSDKNRGKSDN
VPSEEVVKKMKNYWRQLLNAKLITQRKFDNLTKAERGGLSELDKAGFIKRQLVETRQITKHVAQ
ILDSRMNTKYDENDKLIREVKVITLKSKLVSDFRKDFQFYKVREINNYHHAHDAYLNAVVGTALIK
KYPKLESEFVYGDYKVYDVRKMIAKSEQEIGKATAKYFFYSNIMNFFKTEITLANGEIRKRPLIET
NGETGEIVWDKGRDFATVRKVLSMPQVNIVKKTEVQTGGFSKESILPKRNSDKLIARKKDWDP
KKYGGFDSPTVAYSVLVVAKVEKGKSKKLKSVKELLGITIMERSSFEKNPIDFLEAKGYKEVKKD
LIIKLPKYSLFELENGRKRMLASAGELQKGNELALPSKYVNFLYLASHYEKLKGSPEDNEQKQLF
VEQHKHYLDEIIEQISEFSKRVILADANLDKVLSAYNKHRDKPIREQAENIIHLFTLTNLGAPAAFK
YFDTTIDRKRYTSTKEVLDATLIHQSITGLYETRIDLSQLGGDKRPAATKKAGQAKKKKGGGSG
GGSGGGSKDNTVPLKLIALLANGEFHSGEQLGETLGMSRAAINKHIQTLRDWGVDVFTVPGKG
YSLPEPIQLLNAKQILGQLDGGSVAVLPVIDSTNQYLLDRIGELKSGDACIAEYQQAGRGGRGR
KWFSPFGANLYLSMFWRLEQGPAAAIGLSLVIGIVMAEVLRKLGADKVRVKWPNDLYLQDRKL
AGILVELTGKTGDAAQIVIGAGINMAMRRVEESVVNQGWITLQEAGINLDRNTLAAMLIRELRAA
LELFEQEGLAPYLSRWEKLDNFINRPVKLIIGDKEIFGISRGIDKQGALLLEQDGIIKPWMGGEISL
RSAEKAYPYDVPDYA*

#### Supplemental Tables

**Supplemental Table 1 (Related to Figure 2): Limma Analysis of Mass Spectrometry Data**

**Supplemental Table 2 (Related to Figure 3 and 4): Sequences of gRNAs**

| gRNA | Sequence |
|---|---|
| Chr5 Gene Desert #1 | CCAGAAATTAACTGTACCTG |
| Chr5 Gene Desert #2 | GAAGGTCAGACATTAATGTG |
| CD55 #1 | GCTGACTTGGCTTTAGGGGT |
| CD55 #2 | AATGCCCAGCCAGCTTTGGA |
| VEGFA | GACCCCCTCCACCCCGCCTC |
| CD59 #1 | TTCTCAGAACCTGGGCCAGG |
| CD59 #2 | GGCCCAGGTTCTGAGAAGGC |
| c-myc | GCGTCCTGGGAAGGGAGATC |
| Non-coding CD25 | TTGGGCTGGCGTGTTCAGCC |
| Coding CD25 | CACAAGGGTGACAGCCCAGG |
| Non-coding CD55 | TCCAAAGCTGGCTGGGCATT |
| Coding CD55 | AATGCCCAGCCAGCTTTGGA |
| Non-targeting | ACGGAGGCTAAGCGTCGCAA |
| CD25 -48NC | TTGGGCTGGCGTGTTCAGCC |
| CD25 -203NC | GTGGGCTGGGGTTGATGAGA |
| CD25 +17C | CACAAGGGTGACAGCCCAGG |
| CD25 -249C | TTATGGGCGTAGCTGAAGAA |
| CD25 -287C | CACCCTACCTTCAACGGCAG |
| CD25 +0NC | TGGGTCCATCCAGTCTCTAT |

| | |
|---|---|
| CD25 -80C | AGATGAGAGAAGAGAGTGCT |
| CD55 +78C | GGCGCGCCATGACCGTCGCG |
| CD55 +208C | GGGAAGCCCCTGGGCTGGGT |
| CD55 +208NC | GGGAAGCCCCTGGGCTGGGT |
| CD55 +378C | GCTGACTTGGCTTTAGGGGT |
| CD55 +172NC | ACTCACCCCACACGGCCGGC |
| CD55 +731NC | TCCAAAGCTGGCTGGGCATT |
| CD55 +731C | AATGCCCAGCCAGCTTTGGA |
| CD55 +3541C | GCTTCTTGGAAGGCTGAGGC |
| CD55 +3541NC | GCCTCAGCCTTCCAAGAAGC |
| CD55 +8044NC | GCTACAAGGGAGGCTGAGGT |
| CD59 -79NC | CAGGATGCCCTTGCCCTCCC |
| CD59 +245C | TATCTACGAGGAAGGAGAAA |
| CD59 +224NC | AATGAAGCCAGCGTTCGGCT |
| CD59 +143NC | GGTCGAGTGGAAAGCGAGGA |
| CD59 -181C | TTCTCAGAACCTGGGCCAGG |
| CD59 -184NC | GGCCCAGGTTCTGAGAAGGC |
| CD59 -238NC | CCCAGGGAACTGAAAGTTTG |
| CD59 -250NC | AAAGTTTGGGGCGTCCTCCT |
| CD59 +13564C | TCAGACCAAAGGGGTGACTC |
| CD59 +13643C | GTTCCCACTCTACTGGCCCC |
| CD59 +13744NC | CAGTGTGGTAGTACACACTG |

**Supplemental Table 3 (Related to Figure 3): Sequences of ssODN HDR Donors**

| | |
|---|---|
| Chr5 Gene Desert #1 | CATTTAAATCTCTCTACATGAAAAGATAATTGCTCCAGAAATTAACTGTACCTGGAATTCATTCATTTGGTCATGGCTAGTTTCTTATGTAGTGATGATTTGATATCAGAGCTAA |
| Chr5 Gene Desert #2 | TATCAGAGCTAATTAAGATCGGGTCAGAGTTGGTGAAGGTCAGACATTAATGTGTAAGATCATTGAAATGATAATTATAAACAGATTAGGAAGCCACGGTCCTTTATAAGGGGTT |
| CD55 #1 | CCAGCATTTGGGGCTCCTGCTGTGTCGGCCCCCAGCTGACTTGGCTTTAGGGGTAGACGTGGAGGGTTAAAGAGGCCCCGGCTGGGTTTGCGGAGCAGCCAAGCCTGGCAAAATC |
| CD55 #2 | CCTAGGTGACTGTGGCCTTCCCCCAGATGTACCTAATGCCCAGCCAGCTTTGGAAAGCCGTACAAGTTTTCCCGAGGATACTGTAATAACGTACAAATGTGAAGAAAGCTTTGTG |
| VEGFA | GGACGAAAAGTTTCAGTGCGACGCCGCGAGCCCCGACCCCCTCCACCCCGCCACCTGGCGCGGGCTCCGGCCCCTGCCCGCGGCTCGCCGCCGCGTCCACTGTCCGCCGCCGGCC |
| CD59 #1 | TTGAAGGTGCTCATTGGGTCCTGGCCACCCGGCCTTCTCAGAACCTGGGCCAGGATTCTGAGCTCCGCGCGGGGGTGGAGGGAGAGGAGGAGGTTCCTGCCGAGGTGCGGCTGCG |

| | |
|---|---|
| CD59 #2 | TCTCCCTCCACCCCCGCGCGGAGCTCAGCCTCCTGGCCCAGGTTCTGAG AAGGCCTTGTGGCCAGGACCCAATGAGCACCTTCAAAACCCCAGGGAAC TGAAAGTTTGGGGCGTC |
| c-myc | GTGGAAGAGCCGGGCGAGCAGAGCTGCGCTGCGGGCGTCCTGGGAAGG GAGATCCAAAGCGAATAGGGGGCTTCGCCTCTGGCCCAGCCCTCCCGCT GATCCCCCAGCCAGCGGT |

**Supplemental Table 4 (Related to Figure 3): Sequences of Amplicon-NGS PCR Primers**

| | |
|---|---|
| Chr5 Gene Desert gRNA #1 | |
| PCR #1 - Forward Primer | GACTACCTGCCCACATCGTTAC |
| PCR #1 - Reverse Primer | GAGGAAGGAATACACTCTCACC |
| PCR #2 - Forward Primer | gctcttccgatctGACTGTGGAGCCCTGCCTTTG |
| PCR #2 - Reverse Primer | gctcttccgatctCCCCTTATAAAGGACCGTGGC |
| Chr5 Gene Desert gRNA #2 | |
| PCR #1 - Forward Primer | GACTGTGGAGCCCTGCCTTTG |
| PCR #1 - Reverse Primer | CCATCATTGTCCACAGGACAGC |
| PCR #2 - Forward Primer | gctcttccgatctGGGTTCATTCATTTGGTCATGGC |
| PCR #2 - Reverse Primer | gctcttccgatctGGGCTGGAGCTACCATTCTAC |
| CD55 gRNA #1 | |
| PCR #1 - Forward Primer | AGGTCCAAGTCGGTCTCTGAG |
| PCR #1 - Reverse Primer | GGAGACAAAAGCAGAACTGAAGG |
| PCR #2 - Forward Primer | gctcttccgatctCCGCCGTCCTGTGCCTTTAAG |
| PCR #2 - Reverse Primer | gctcttccgatctCCACACGGCTGGACTCTGTC |
| CD55 gRNA #2 | |
| PCR #1 - Forward Primer | CCACTCTCGACAGAGTCCAGC |
| PCR #1 - Reverse Primer | GTGACGTGCCAACAGGGTATAC |
| PCR #2 - Forward Primer | gctcttccgatctGCAACTGTGAGGACACTTGATAG |
| PCR #2 - Reverse Primer | gctcttccgatctCAATATCTGACCATTGACTGCCC |
| VEGFA | |
| PCR #1 - Forward Primer | GAGGTAGCAAGAGCTCCAGAGAG |
| PCR #1 - Reverse Primer | CTGGAGCACTGTCTGCGCACA |
| PCR #2 - Forward Primer | gctcttccgatctTGACGGACAGACAGACAGACAC |
| PCR #2 - Reverse Primer | gctcttccgatctGGCCCGAGCTAGCACTTCTC |
| CD59 gRNA #1 | |
| PCR #1 - Forward Primer | GTAGGAAGCAGCTTCAGACTGC |
| PCR #1 - Reverse Primer | CTCGGCTCGGCTCACCCAAAC |
| PCR #2 - Forward Primer | gctcttccgatctGCAAATCCGAGGAGGACGCC |
| PCR #2 - Reverse Primer | gctcttccgatctCATTCTTTCGCTCCAGCCCGCA |
| CD59 gRNA #2 | |

| | |
|---|---|
| PCR #1 - Forward Primer | GAAGCAGCTTCAGACTGCAGC |
| PCR #1 - Reverse Primer | GCCTTCGGGCCTTCTTACCTG |
| PCR #2 - Forward Primer | gctcttccgatctATGTCCCATAGCAAATCCGAGG |
| PCR #2 - Reverse Primer | gctcttccgatctCCCGCATTCTTTCGCTCCAGC |
| c-myc gRNA | |
| PCR #1 - Forward Primer | CCCTTTATAATGCGAGGGTCTG |
| PCR #1 - Reverse Primer | AATCCAGCGTCTAAGCAGCTGC |
| PCR #2 - Forward Primer | gctcttccgatctGGGCTTTATCTAACTCGCTGTAG |
| PCR #2 - Reverse Primer | gctcttccgatctTGCTATGGGCAAAGTTTCGTGG |

**Supplemental Table 5 (Related to Figure 4): Sequences of qPCR Primers for Chromatin Immunoprecipitation**

| | |
|---|---|
| Template CD25 gRNA | |
| Upstream of Protospacer - Forward Primer | CTTCTCATCAACCCCAGCCC |
| Upstream of Protospacer - Reverse Primer | CCTAGCACTCTCTTCTCTCATCTC |
| Inclusive of Protospacer - Forward Primer | GAGATGAGAGAAGAGAGTGCTAG |
| Inclusive of Protospacer - Reverse Primer | ACCCTTGTGGGTCCATCCAG |
| Non-Template CD25 gRNA | |
| Upstream of Protospacer - Forward Primer | CTTCCCATCCCACATCCTCC |
| Upstream of Protospacer - Reverse Primer | CCCACATCAGCAGGTATGAATCC |
| Inclusive of Protospacer - Forward Primer | GAGAGCAACTCCTGACTCCG |
| Inclusive of Protospacer - Reverse Primer | CTTTCTCTGCAGAAGGCCCA |
| CD55 gRNAs | |
| Upstream of Protospacer - Forward Primer | CCACTCTCGACAGAGTCCAG |
| Upstream of Protospacer - Reverse Primer | CAAGTGTCCTCACAGTTGCTG |
| Inclusive of Protospacer - Forward Primer | CCTAGGTGACTGTGGCCTTC |
| Inclusive of Protospacer - Reverse Primer | GGCAGATCACTGAGTCCTTCTC |
| Reference (c-myc) | |
| Forward Primer | GCCGCATCCACGAAACTTTG |
| Reverse Primer | GCAAGGAGAGCCTTTCAGAGAAG |
